# Differentiation of *Xanthomonas oryzae* pv. *oryzae in vitro* and during rice leaf infection

**DOI:** 10.1101/2025.10.12.680524

**Authors:** Laura Redzich, Zongyi Ma, Andrea Restrepo-Escobar, Miriam Bäumers, Van Schepler-Luu, Eliza P.I. Loo, Wolf B. Frommer

**Author notes:** Corresponding author: Wolf B. Frommer.

## Abstract

**Highlights:**

– Xoo produces filamentous morphology, which is transient and yields pleomorphic progenies *in vitro*
– *In planta*, initial attachment of rod-shaped Xoo is detected at xylem pits
– The Xoo infection front migrates basipetally in the vascular bundle and progresses laterally from major to minor veins via transverse veins
– Xoo breaks out of the xylem vessels and enter the neighboring xylem parenchyma
– Xoo assumes filamentous morphology that can traverse from the xylem across the bundle sheath into mesophyll tissue
– Mobility in xylem vessels depends predominantly on rod-shaped Xoo, while infection of mesophyll tissue at later stages appears to be linked to filamentous morphology

**Summary:** *Xanthomonas oryzae* pv. *oryzae* (Xoo) is classified as a xylem pathogen responsible for bacterial blight of rice causing substantial yield losses in Asia and Africa. Xoo virulence depends on the ability to trigger SWEET sucrose efflux transporters in the xylem parenchyma (XP) by injection of transcription activation like effectors (TALe) into host cells, likely to access host-derived sucrose. To establish infection, Xoo must overcome physical barriers, immune responses and the hydraulic xylem flow. To gain insights into the colonization process, we used translational *SWEET11a-GUS* reporter lines, scanning electron microscopy, and confocal laser scanning microscopy of Xoo tagged with a fluorescent protein. We found that Xoo can differentiate *in vitro* into filamentous forms. We mapped the infection route of Xoo along the vasculature, identified distinct spatiotemporal phases of Xoo colonization marked by rod-shaped and, notably, filamentous Xoo cells. Rod-shaped Xoo were found to attach to xylem pits during basipetal progression of the infection. Notably, we found that at later infection stages, Xoo could enter the XP. Strikingly, Xoo adopted a filamentous phenotype that traversed bundle sheath cells and entered mesophyll cells. Chlorosis and necrosis of leaves is thus likely not just due to blockage of xylem flow, but to direct tissue damage. Filamentation had been reported as important for virulence of human pathogens e.g. *Yersinia pestis*, uropathogenic *E. coli* and *Shigella* and had been associated to sugar utilization in *Bacillus subtilis*. We thus hypothesize that Xoo differentiation during host colonization is critical for virulence.

**Graphical abstract:** 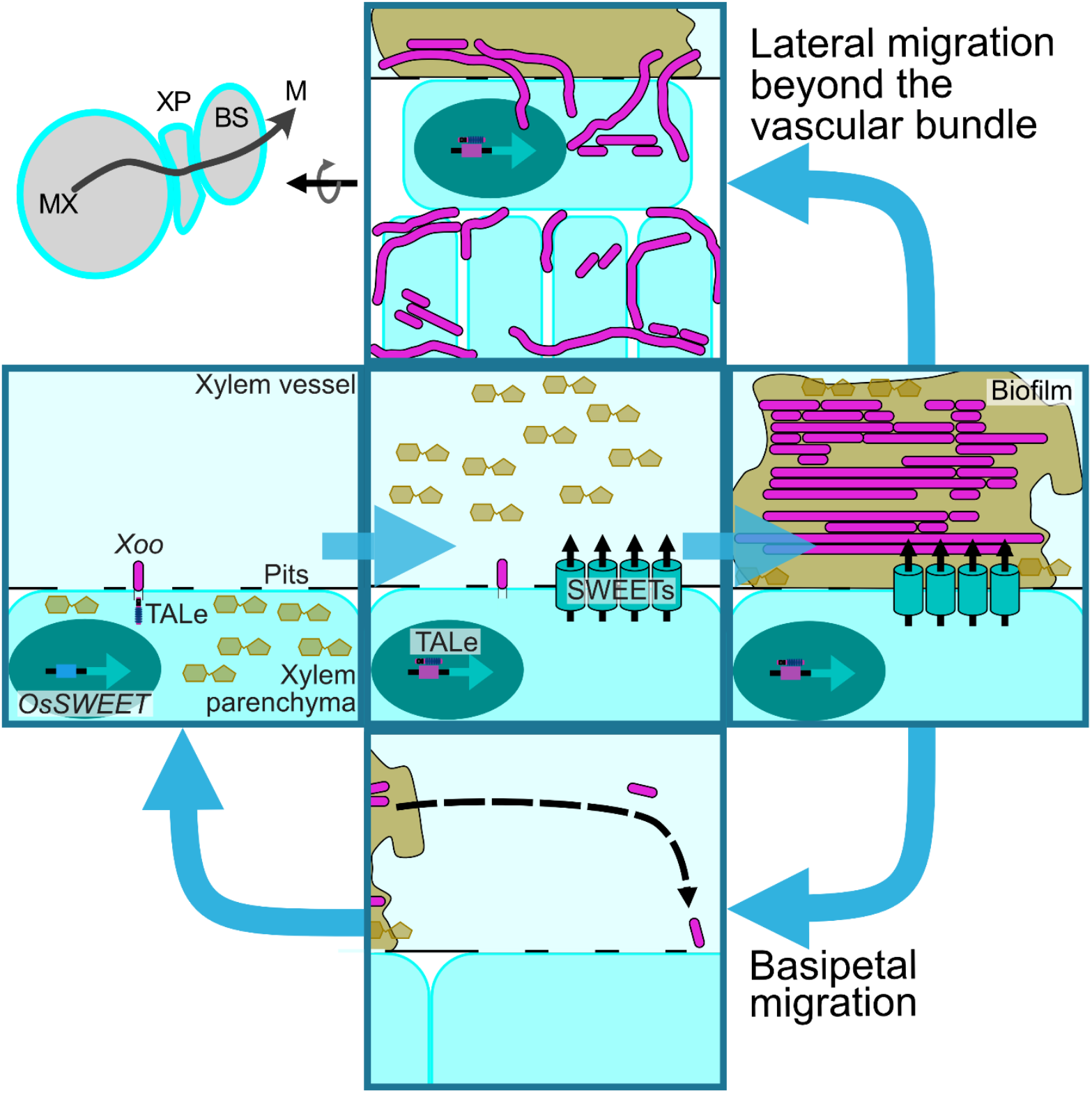

## Introduction

Pests and pathogens cause massive yield losses worldwide and impact food security (1). Among the different organism that cause losses, phytopathogenic bacteria are a major factor (2). 5-20 % of annual yield losses are reported for the causal agent of bacterial blight, *Xanthomonas oryzae* pv. *oryzae* (Xoo) in Asia and Africa (1,3). The gram-negative bacterium Xoo enters rice (*Oryzae sativa*) leaves through hydathodes or wounds, enters xylem vessels and subsequently multiplies in the xylem (4,5). A critical aspect of Xoo virulence is its dependence on host-derived sucrose (6–8). To access sucrose, Xoo utilizes a type III secretion system (T3SS) to deliver transcription activator-like effectors (TALe) to adjacent xylem parenchyma cells (XP) that are translocated into nuclei and induce *SWEET* sucrose uniporter genes (6,8,9). The induction of *SWEET* uniporters likely causes sucrose efflux from XP cells, where it becomes available to Xoo through bacterial transporters and sucrose hydrolases encoded by the sucrose utilization cistron *sux* (10).

Xoo has to overcome multiple challenges during host infection, e.g., acquire a full complement of all essential nutrients, migrate against the xylem stream, resist the forces of the flow, and evade host immune system responses. Extracellular sucrose acts as chemoattractant for Xoo, induces the production of quorum sensing molecules and taken up and metabolized by the components of the sux gene cluster of Xoo (10–12).

Rather than acting as solitary units, bacterial cells can form structured communities that function in unison (Ng et al., 2009). Bacteria have been shown to change cell shape or become multicellular, either as chains (i.e., chaining) or as filaments with or without septation. Septation requires treadmilling of FtsZ to produce septal peptidoglycans (14). In the absence of FtsZ and peptidoglucan synthetic activity, septate or chained filaments are produced without cell separation. *In vitro*, conditional filamentation can be triggered by a variety of factors, e.g., antibiotics, anaerobiosis, high osmolality, pH shifts, temperature shock, or UV. Changes in nutrient supply can also induce filamentation, e.g., thymine deprivation, low or high Mg^2+^ l, high PO_4_^3-^, and limiting or excessive amino acids (15–18). Bacterial filamentation has been described as an adaptive strategy to enhance resilience and survival under unfavorable conditions by minimizing cell division (19–21), and it is important for biofilm formation and bacterial dispersal (22,23). Filamentation was reported as a defense mechanism to flagellate grazing (24,25).

Filaments of uropathogenic *Escherichia coli* were associated with higher tolerance to host defenses (26,27). It has been suggested that due to their size, filaments could be less prone to internalization by macrophages (23,28). Several studies on the effects of antibiotics on bacterial morphology have reported filamentation in pathogens (29,30), such as the effect of chelerythrine on Xoo (31). Filamentation of human pathogens has been linked to important roles in virulence, e.g., for *Yersinia pestis* or *Vibrio cholerae* (17,32,33). Plant pathogens, e.g., *Xylella fastidiosa*, were reported to filament *in vitro* (18,22,34–38) but their role during host colonization *in vivo*, except for *Dickeya dandtii* (39) has remained elusive and only a few pathogens have been studied *in vivo* in context of host colonization (40,41).

Here, we found that Xoo colonies on solid agar surfaces underwent a pleomorphic shift leading to filamentous forms. In liquid cultures high salinity and elevated cell density induce conditional filamentation of Xoo *in vitro*. To explore whether Xoo can also filament during the infection of rice leaves, we tracked the infection process using imaging of reporter activity of translational SWEET11a-GUS reporter lines. Accumulation of diX Indigo (product of GUS) served as a proxy for the stage of successful injection of TALes into XP, enabling spatiotemporal mapping of Xoo during the infection process. Xoo infection appeared non-continuous within main veins at the infection front and progressed into minor and lateral veins. Data on the infection route were then used to guide high-resolution imaging by confocal laser scanning microscopy (CSLM) and scanning electron microscopy (SEM). Rice leaf autofluorescence was overcome by tracking Xoo expressing the large Stokes shift fluorescent protein LSSmApple. At the infection front, rod-shaped Xoo were arranged perpendicular to xylem vessel pits. Strikingly, we discovered a previously unreported pleomorphic shift from rod-shaped to filamentous Xoo cells during host infection *in planta*. Time-lapse imaging revealed that filamentation was transient and asymmetric cell division yielded pleomorphic progenies. Notably, at later stages of the infection, Xoo was able to enter the XP, likely by hydrolysis of host pit cell walls. Filamentous Xoo spread laterally beyond the vascular bundle (XP and bundle sheath cells) even to the mesophyll, consistent with the observed disease symptoms that lead to chlorosis in the mesophyll and necrosis.

## Results

### Phenotypic heterogeneity of *in vitro* colonies

Morphological plasticity is common among pathogens (22,26,39,42). To test whether Xoo colonies exhibit phenotypic heterogeneity on the macroscopic scale, colonies were grown for six days on solidified NBS agar plates. To quantify cross-colony variability, images of colonies were analyzed using hue, saturation and brightness (HSB) criteria (Figures 1A, S1). The HSB color-space approach enabled pixel-level discrimination of structural and optical features across the colony surface. The analysis focused on grayscale variation of individual pixels, with the assumption that variation in brightness and saturation could reflect underlying differences in colony density and cell morphology. Based on the analysis, colonies were reproducibly segmented into three concentric growth zones, each characterized by distinct HSB profiles (Figure S1). To evaluate the phenotypic appearance of Xoo across the colony center (1), mid-zone (2) and peripheral edge (3), Xoo was visualized by CSLM (Figure 1B). Rod-shaped and filamentous Xoo cells appeared (defined as: < 5 µm - rod-shaped, > 5 µm - intermediate and filamentous) throughout all three zones. However, the colony center (1) contained the highest proportion of rod-shaped cells, while the peripheral edge (3) contained the largest number of filamentous cells (Figure 1C). The mid-zone (2) exhibited an intermediate distribution. The spatial pattern indicates that Xoo adopts morphologies in response to local microenvironments in the colony (Figure 1D).

**Figure 1.**
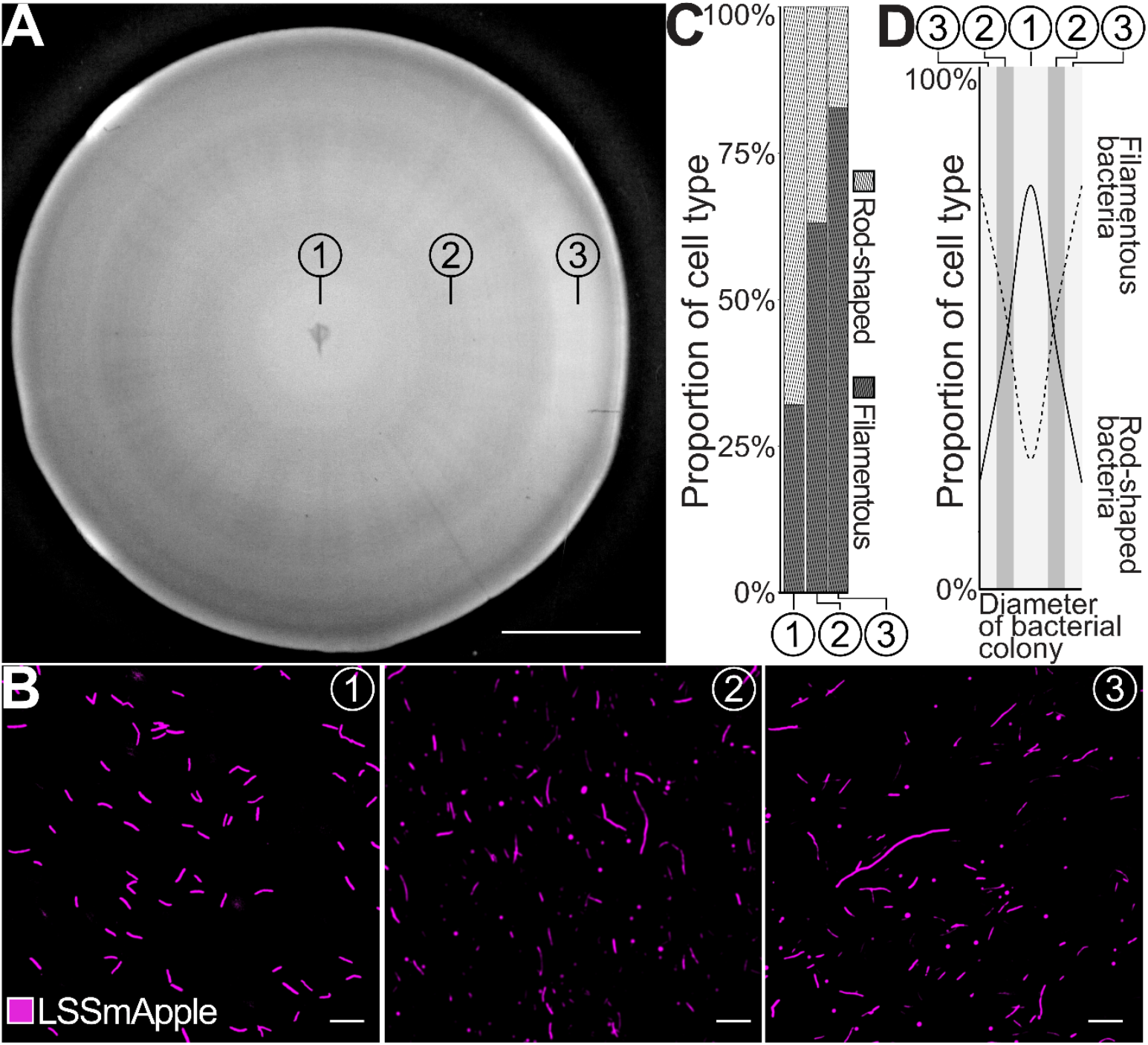
Pleomorphism and spatial zonation in *Xanthomonas oryzae* pv. *oryzae* colonies on solid media. **(A)**. Colony-scale imaging of six-day old colonies of PXO99^A^_LSSmApple_ revealed spatial zonation into three concentric regions: colony center (1), mid-zone (2) and peripheral edge (3). Scale bar: 5 mm. (**B)**. Pleomorphism of PXO99^A^_LSSmApple_ was discovered by high resolution microscopy of individual concentric zones. Images were taken with ZEISS LSM880 Airyscan (excitation: 488 nm, detection range: 605-615 nm) Scale bar: 10 µm. **(C)**. For each concentric zone, cell lengths of 100 bacteria were quantified and grouped into rod-shaped (< 5 µm) or filamentous (cells > 5 µm, encompassing intermediate (5-10 µm) and filamentous morphologies (>10 µm)). Quantification of cell length was repeated independently three times on different colonies with comparable results **(D)**. The represented bacterial morphologies across PXO99^A^_LSSmApple_ colonies were illustrated based on (B). Related to Figure S1. HSB color space segmentation of bacterial colonies.

### Morphological plasticity of individual Xoo

To examine whether Xoo filamentation can be triggered independent of a surface-attached colony, we exposed bacteria to two conditions in liquid culture: 500 mM NaCl (hereafter ‘high salinity’) and high cell density (OD_600_ 3, hereafter ‘high OD_600_’). Bacterial cell length was assessed using CLSM (Figure 2A). High salinity caused the largest heterogeneity in cell length with a mean average of 9.6 ± 6.9 µm. Although significantly shorter, bacteria from high OD_600_ also exhibited elongated forms, with a mean length of 4.2 ± 3.1 µm. In comparison, control cultures at OD_600_ 0.4 displayed a narrow distribution of cell lengths (mean 2.4 ± 0.6 µm) significantly different from high salinity and high OD_600_ conditions. The results indicate that Xoo filamentation can be induced *ex planta* by specific environmental cues. During host colonization, bacteria need to maintain microcolonies and need to disperse to gain access to nutrients and to spread; filaments could serve as intermediate stages. Some multinucleate filaments were shown to rapidly divide into rod-shaped cells after release from sublethal conditions (21,43,44). To explore whether reproductive activity of Xoo cells depends on cell length and growth history, we evaluated the number of colony forming units (CFU) from different morphological states. Subpopulations of <5 µm (rod-shaped), 5–10 µm (intermediate), and >10 µm (filamentous) cells were enriched from high OD_600_ and high salinity populations, washed, set to OD_600_ 0.1 and diluted 10^-6^ fold. CFU were quantified after four days of growth in dark conditions at 28°C. Surprisingly, founder cells from high salinity and high OD_600_ cultures showed differences in CFU depending on their cell length (Figure 2B). While CFU increased with increasing cell length for founder cells from high salinity cultures, CFU decreased with increasing cell length for founder cells originating from high OD_600_ cultures. Filamentous cells originating from high salinity conditions formed significantly more CFU relative to filamentous cells from high OD_600_ cultures. In contrast, rod-shaped cells from high OD_600_ cultures formed most colonies, intimating that rod-shaped cells were more actively reproducing within the high OD_600_ community. Intermediates originating from high salinity cultures exhibited a higher CFU compared to intermediates from high OD_600_ cultures, further supporting the observation regarding differences in reproduction activity for rod-shaped and filamentous cells between communities from different growth conditions. Our observations imply that subpopulation-specific traits differentiate with growth history.

**Figure 2.**
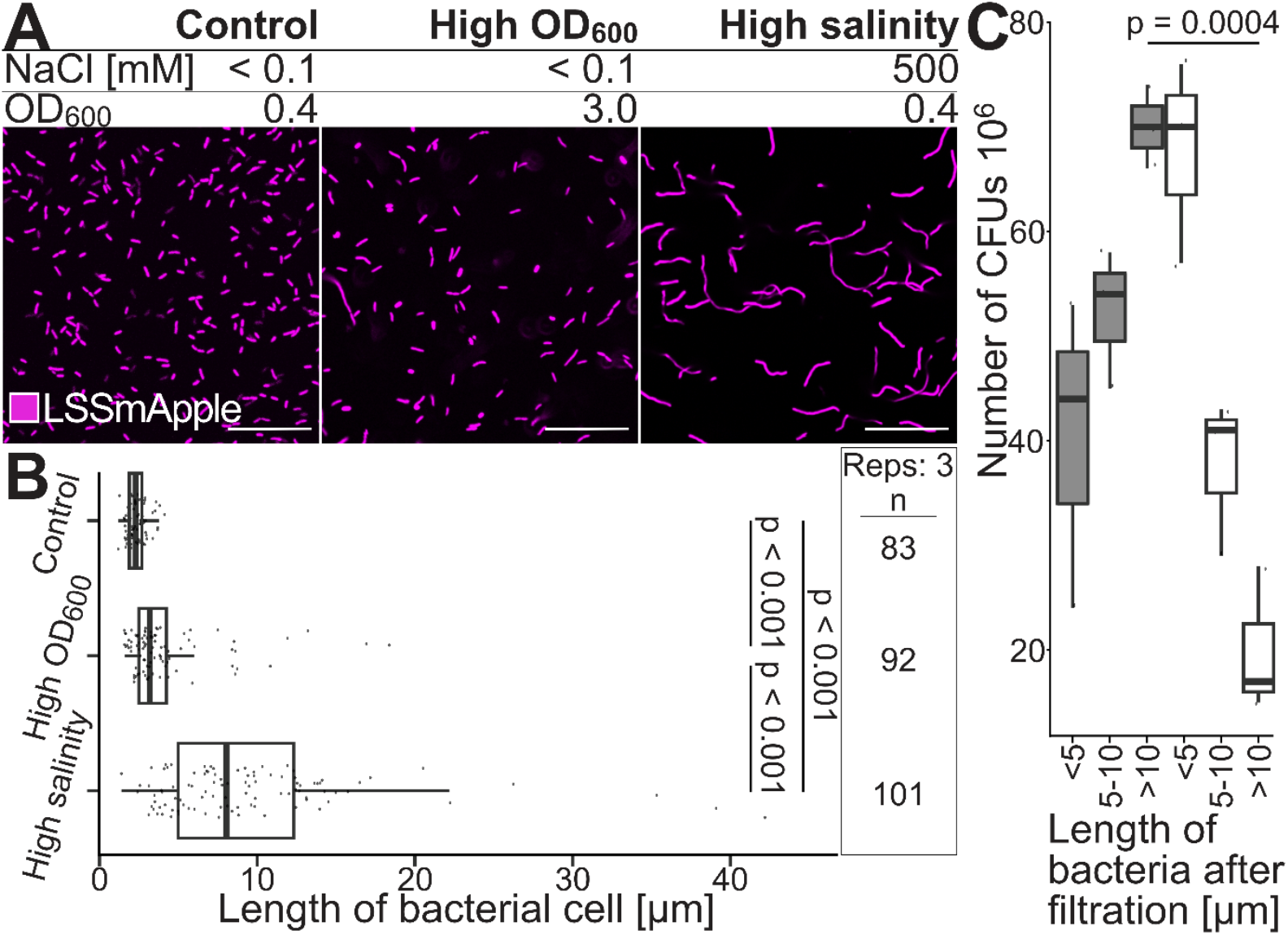
Conditional filamentation and viability of *Xanthomonas oryzae* pv. *oryzae* in response to different growth conditions in liquid culture. **(A)**. PXO99^A^_LSSmApple_ was cultivated in liquid cultures (150 rpm, 28 °C, dark conditions) supplemented with 500 mM NaCl (high salinity) or to OD_600_ 3 (high OD_600_) and observed by confocal laser scanning microscopy (ZEISS LSM880; excitation: 488 nm, detection range: 605-615 nm). (**B)**. Cell length of PXO99^A^_LSSmApple_ cultivated to OD_600_ of 0.4 or OD_600_ of 3 in <1 mM of NaCl, and OD_600_ 0.4 in 500 mM NaCl were measured with the Segmented Line Tool in ImageJ. Data processing was performed in R. Repeated independently three times with comparable results. Scale bar: 10 µm. **(C)**. High OD_600_ (white) and high salinity populations (grey) were divided by length into rod-shaped (< 5 µm), intermediate (5-10 µm) and filamentous (> 10 µm) subpopulations. Subpopulations were diluted by 10^6^ and plated in triplicates onto petri dishes with 20 mL NBSA media supplemented with 100 µg/mL Kanamycin. Colony forming units were quantified after three days at 28 °C, dark conditions. Data processing was performed in R. CFU assay was independently repeated three times with comparable results.

### Filament differentiation

To explore the potential role of subpopulation-specific traits, differentiation of filaments was observed by time lapse imaging of filamentous Xoo using transmission light microscopy and CSLM (Figure 3). Filaments initially underwent further elongation, extending from 13.4 ± 3.2 µm to 28.3 ± 5.1µm after 3 hours. After elongation, elongated filaments showed septum formation as soon as 4 hours after transferal to imaging chamber, with septa positioned at irregular intervals along the cell body (indicated by arrows, Figure 3 A, B, S2,3). The septation events ultimately led to asymmetric division and re-establishment of a morphologically heterogeneous population of filamentous, intermediate and rod-shaped cells. After 17 hours, the culture exhibited a dense mixed population, demonstrating that morphologically diverse bacterial populations can be reconstituted from filamentous cells. Different from the unipolar growth of *Corynebacteriu matruchotii* or *Leptrothis cholodnii*, Xoo showed a bipolar elongation phase and irregular septation, followed by asymmetric divisions (Figure 3C). Phenotypic heterogeneity was reconstituted from filamentous parental cells, indicating that pleomorphism is likely connected to intracommunity communication and potentially to subpopulation specific traits.

**Figure 3.**
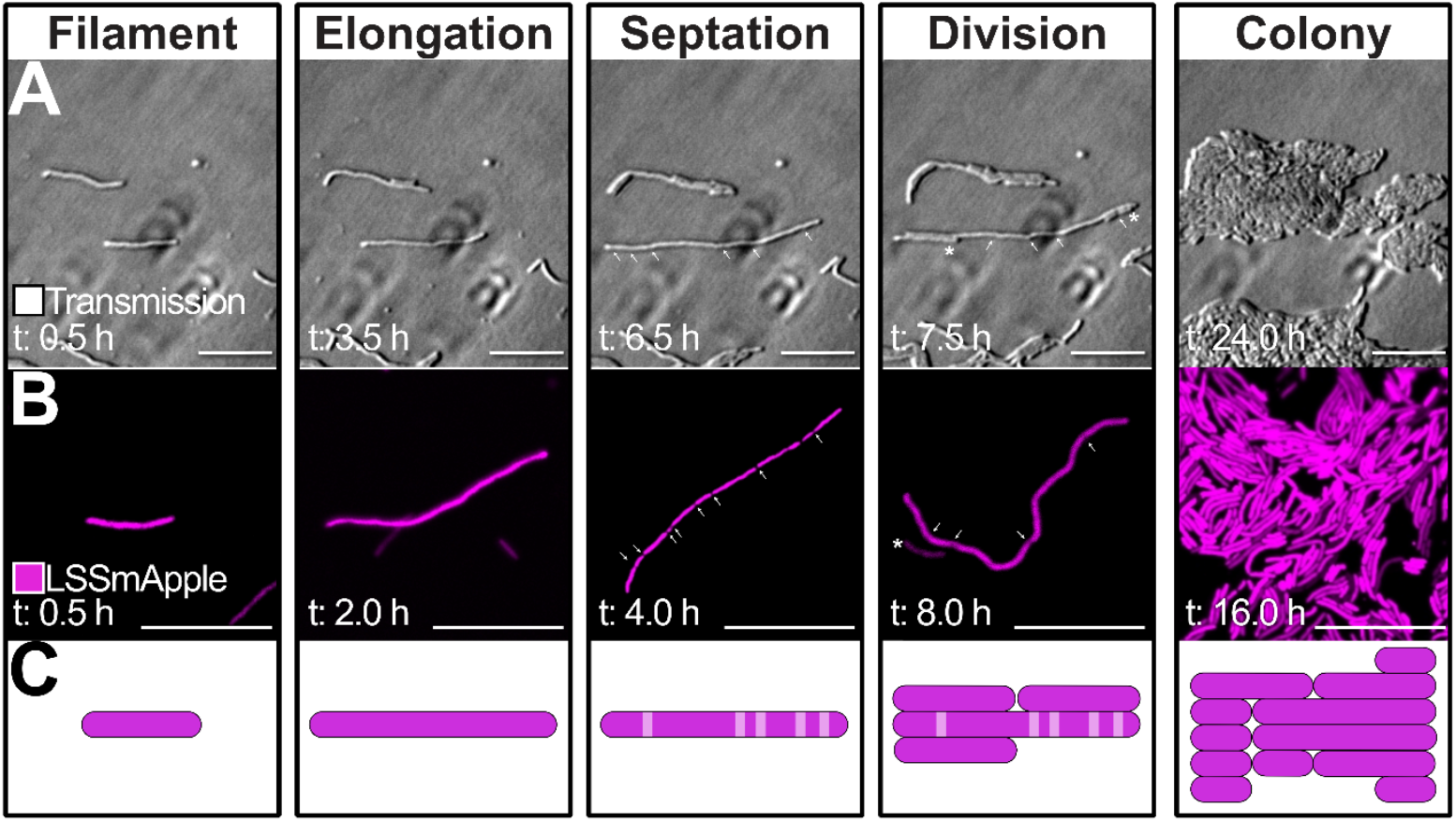
Filamentous *Xanthomonas oryzae* pv. *oryzae* initiate the formation of new colonies that display phenotypic heterogeneity following a developmental program. Filamentous PXO99^A^_LSSmApple_ separated by filtration from cultures grown in high salinity (OD_600_ 0.4, 500 mM NaCl) and placed onto NBSA media within the aerated imaging set-up (Figure S4). **(A)**. Individual filaments were observed for 24 hours using transmission light microscopy (bright field, Nikon Ti Eclipse PFS). Representative images of bacteria at different time points after release from high salinity conditions are shown. Z-stack images were obtained every 30 minutes for 24 hours (Figure S5). Experiments were repeated independently three times with comparable results. Scale bar: 10 µm. (**B)**. PXO99^A^_LSSmApple_ were observed by confocal laser scanning microscopy (ZEISS LSM880; excitation: 488 nm, detection range: 605-615 nm). Representative images at different time points after release of PXO99^A^_LSSmApple_ from high salinity conditions are shown. Experiments were repeated independently three times with comparable results. Scale bar: 10 µm. **(C)**. Cartoon of developmental program followed by filamentous PXO99^A^_LSSmApple_ derived from high salinity conditions after placement onto fresh media. Filaments were observed to elongate in a bipolar fashion. Asymmetric septation was succeeded by division into pleomorphic colonies. Related to Figures: Figure S2. Assembly of an agar-based aerated imaging chamber. Figure S3. Time-lapse imaging of filamentous *Xanthomonas oryzae* pv. *oryzae*.

### Reporter-based mapping of the infection route

We used a combination of bacterial (fluorescent Xoo) and host reporters (GUS reporter lines) to trace the infection. For effective virulence, Xoo must induce at least one of the clade III *SWEET* genes. We used translational SWEET11a-GUS reporter lines to map a specific stage of the infection route of Xoo, i.e., successful attachment to xylem walls, successful formation of the T3SS and injection of the TALe (6,45). Accumulation of the GUS product diX Indigo served as proxy indicator for this infection stage (Figure 4). As expected, diX Indigo accumulated during infection with PXO99^A^_LSSmApple_, which harbors the *SWEET11a* inducing TALe PthXo1, but was absent during infection with the ME2 control strain lacking cognate TALe able to target a clade III *SWEET* promoter (Figure 4A). From the clipping site, progressive basipetal infection was detected; initially in main veins, subsequently in transverse and minor veins (Figure 4A). Using emission scans of rice leaves we identified the large Stokes shift protein LSSmApple to be suited to overcome rice tissue autofluorescence at excitation with 488 nm (Figure S4). The *SWEET11a* inducing Xoo strain PXO99^A^ was transformed with a LSSmApple cassette (PXO99^A^_LSSmApple_). PXO99^A^_LSSmApple_ was then used to monitor the bacteria in the rice leaf at different stages by fluorescence microscopy (Figures 4B). Overall, the patterns were highly similar, however the GUS reporter assay had higher sensitivity also for identifying the bacterial infection front (Figure 4C).

**Figure 4.**
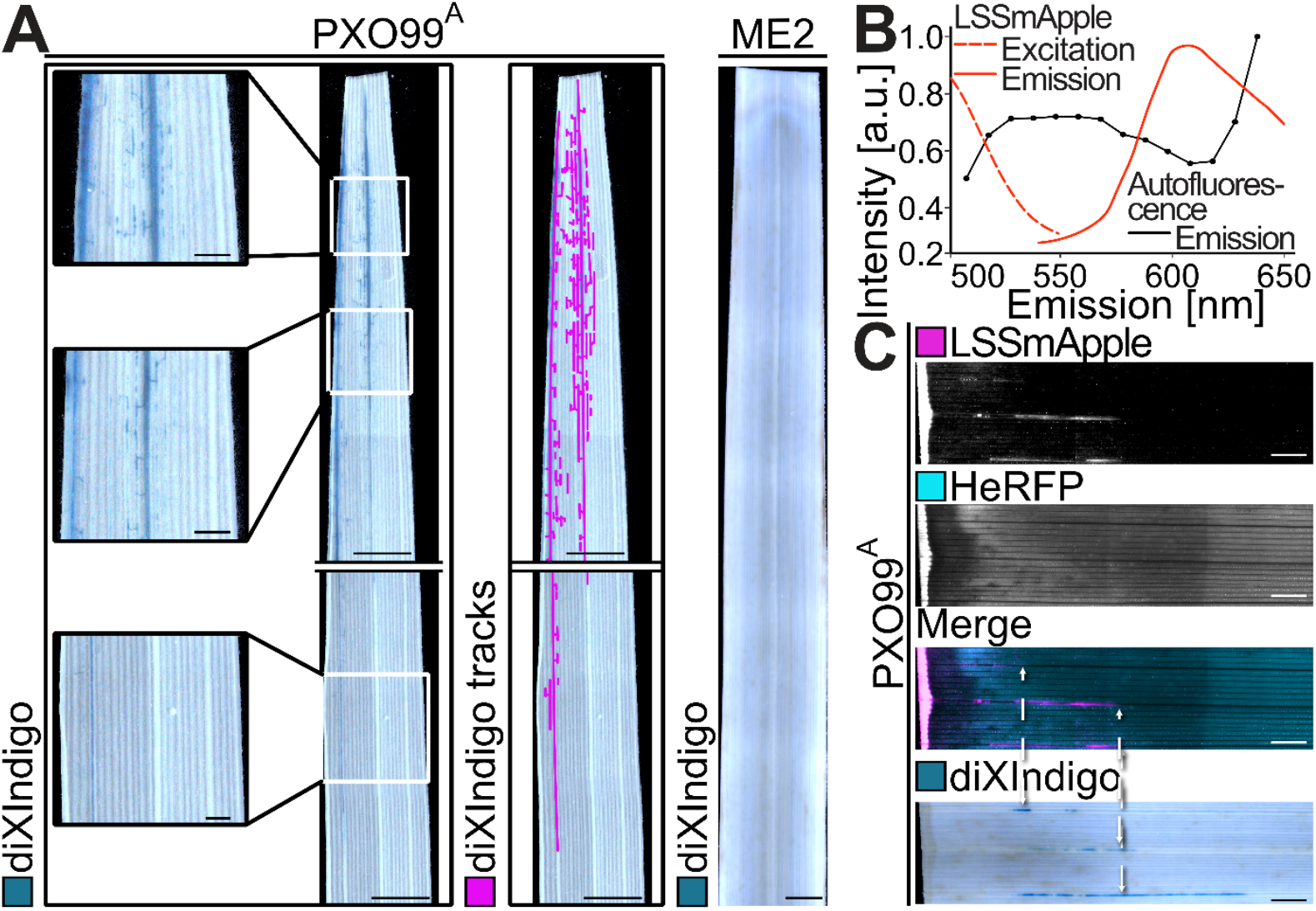
Mapping of the infection process using SWEET11a-GUS reporter lines and Xoo expressing a fluorescent protein PXO99^A^_LSSmApple_. **(A)**. diX Indigo accumulation at 4 days after leaf clipping infection (72) with PXO99^A^ and ME2 observed with the ZEISS Axiozoom.V16. Representative images from three leaves per treatment from three independent experiments performed with two independent pSWEET11a:SWEET11a-GUSplus lines (transformation event #8 and #10)(6). Enlarged sections of the top, mid and bottom zone of the leaf indicated by squares (scale bar: 2 mm). diX Indigo served as indicator for bacterial presence and was tracked in ImageJ (pink, scale bar: 1 cm). ME2 served as negative control (scale bar: 1 cm). **(B)**. Emission scan of rice leaves at 488 nm excitation identified LSSmApple as suitable, monochromatically excitable protein to circumvent autofluorescence of rice leaves. **(C)**. Leaves infected with PXO99^A^_LSSmApple_ were observed 3 dpi with fluorescence microscopy (Zeiss Axiozoom.V16). Subsequently, GUS histochemistry was applied to the same samples. Front of infection depicted by diX Indigo and LSSmApple are indicated by dashed lines. Experiment was repeated three times independently on five leaves each with comparable results. Scale bar: 1 cm. Related Figures: Figure S4. Autofluorescence scans of rice leaves. Figure S5. Selection of infection front for SEM.

### Distinct cell morphologies during host colonization

SEM and CLSM were employed to investigate Xoo morphologies at different infection stages at single cell resolution by monitoring PXO99^A^_LSSmApple_ fluorescence and SEM. Guided by spatiotemporal patterns of Xoo infection established through translational SWEET11a*-*GUS reporter line assays (Figure 4A), we analyzed fixed sections of infected rice leaves corresponding to the infection front and the subsequent progression of the infection (Figure S5). In tissue sections at the bacterial front of infection, rod-shaped PXO99^A^_LSSmApple_ were exclusively observed within the xylem vessel (Figure 5). Remarkably, the majority of PXO99^A^_LSSmApple_ cells located within or in contact with xylem vessel pits (Figures 5A,B). The pit localization is likely the optimal site for T3SS injection due to the thinned cell walls and the site of SWEET-based sucrose efflux. Within the group of PXO99^A^_LSSmApple_ located in xylem vessel pits, 45 % of cells appeared to be orientated perpendicular to the xylem vessel wall (on average in twelve biological replicates; Figures 5C,D), either caused by polar adhesins or polar T3SS attachment. The region spanning 3–5 cm from the leaf tip at six days post infection (dpi) were chosen as representative of an established infection stage of vascular colonization (Figures 6A, S4). At the 6-dpi stage, PXO99^A^_LSSmApple_ displayed striking morphological polymorphism, with a prominent filamentous form not previously documented for Xoo *in planta*. SEM revealed high cell density accumulation of filamentous cells within xylem vessels, often filling substantial portions of the lumen (Figure 6 B,C). CLSM also detected filamentous cells in the metaxylem of major and minor veins (Figures 7, S7-9).

**Figure 5.**
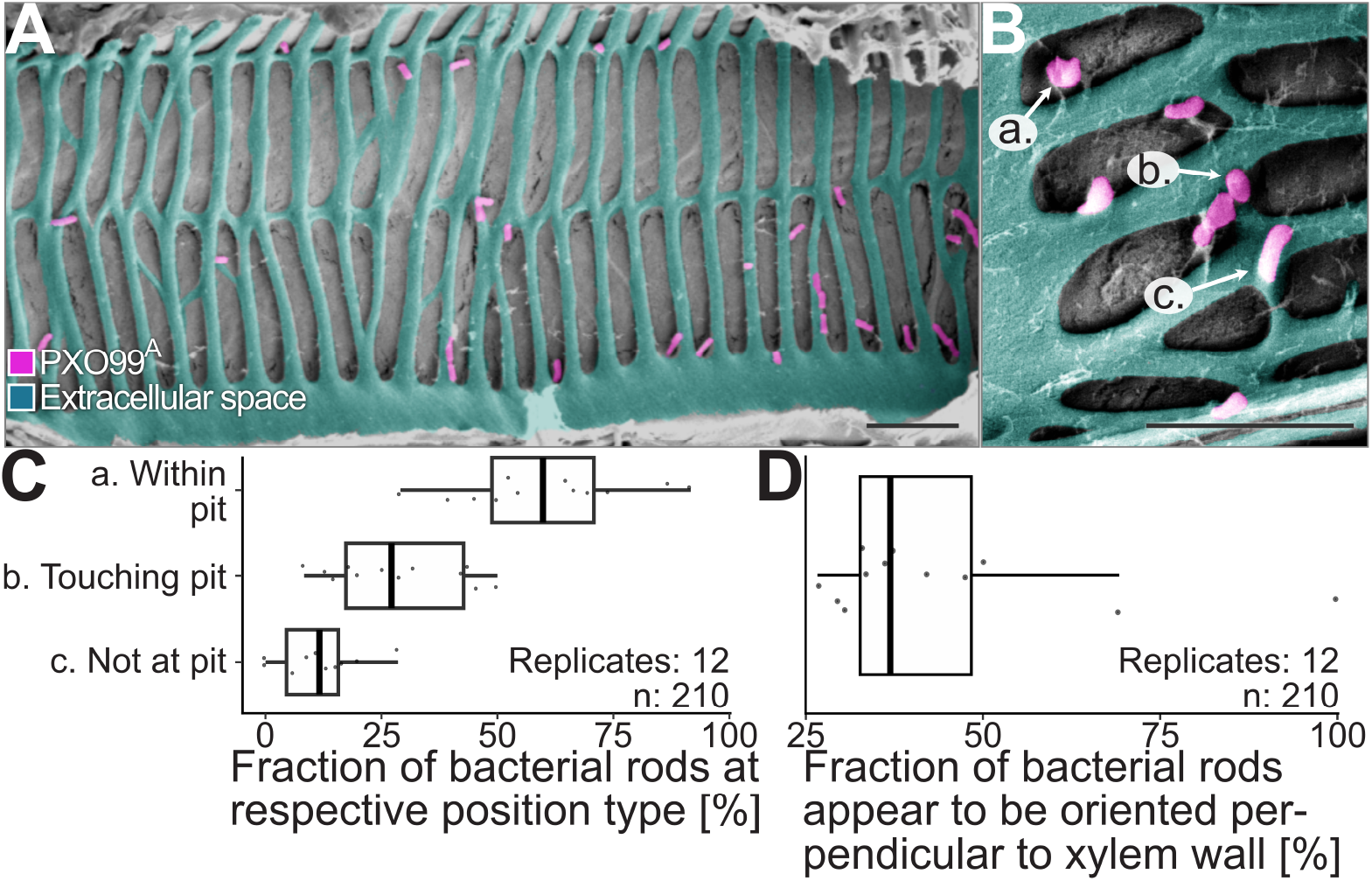
Rod-shaped *Xanthomonas oryzae* pv. *oryzae* locate predominantly to xylem pits near the infection front. **A)**. Samples were taken from the infection front (Figure S6) and processed for scanning electron microscopy. Images were taken scale bar: 5 µm. Post-processing was performed in GIMP (v2.10.24; pink: bacteria, turquoise: secondary xylem vessel wall). **(B)**. Location of Xoo was categorized as within pit (a.), touching pit (b.) and not at pit (c.). Representative images of categories a., b., and c. are presented. Post-processing was performed in GIMP (v2.10.24; pink: bacteria, turquoise: secondary xylem vessel wall). Scale bar: 5 µm. **(C)**. Location of Xoo at the infection front from twelve images from five leaves. Experiment was repeated independently three times with comparable results. Data processing in R. **(D)**. Appearance of Xoo to be orientated perpendicular within the 210 bacterial cells analyzed from (C). Experiment was repeated independently three times with comparable results. Data processing performed in R. Related Figures: Figure S5. Selection of infection front for SEM.

**Figure 6.**
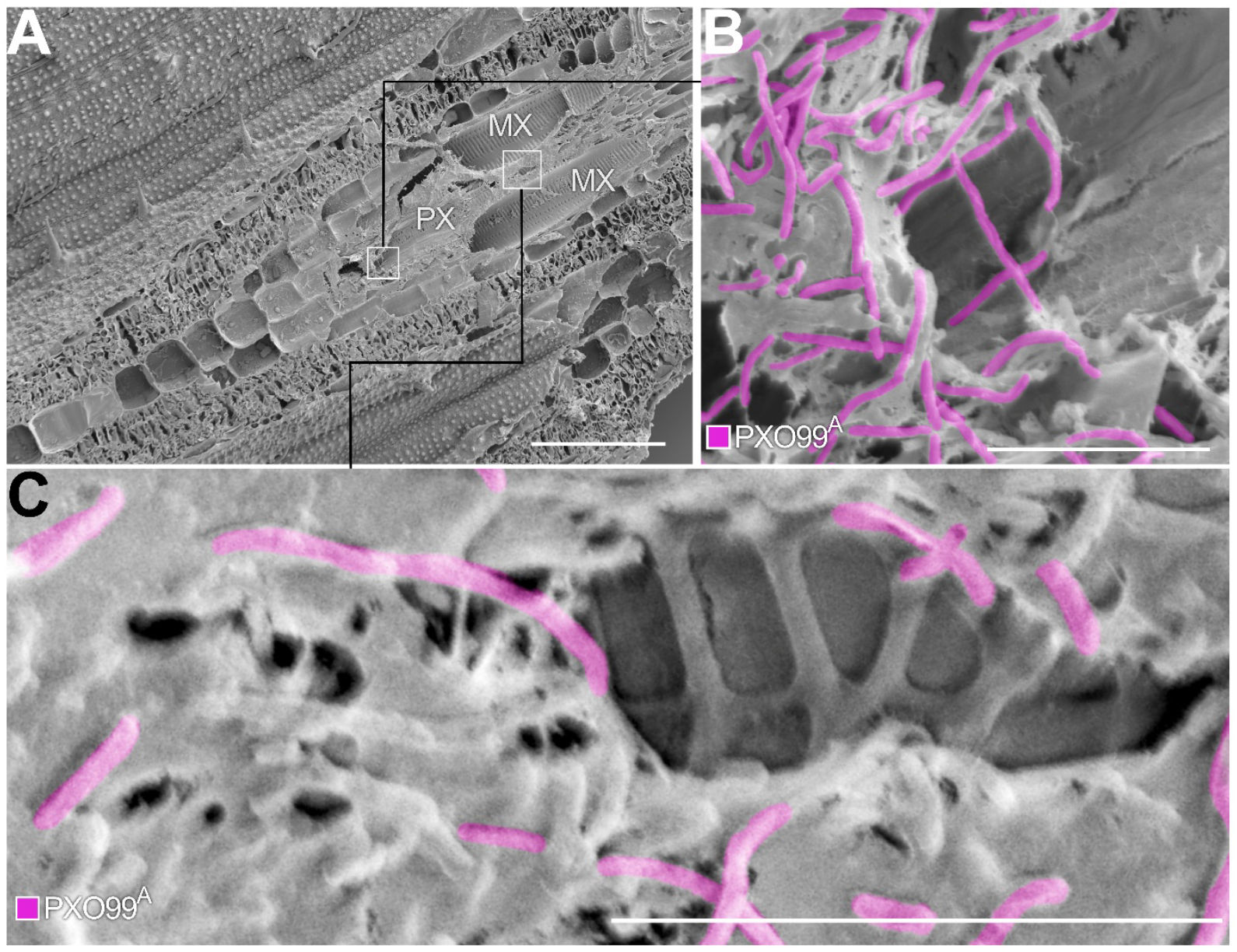
Filamentation of *Xanthomonas oryzae* pv. *oryzae* during host colonization is detected apoplasmically in xylem vessels and intracellularly in xylem parenchyma and adjacent cells. Samples corresponding to an advanced stage of colonization (3-5 cm from the leaf tip at 6 dpi, Figure 4) were fixed and subjected to SEM. Experiment was performed three times independently with comparable results. **(A)**. Overview of infected rice leaf section by SEM, scale bar: 100 µm. Squares indicate regions were magnified images (B, C) were taken. Protoxylem (PX; magnification (B).) is located adaxially to metaxylem vessels (MX, magnification (C). **(B)**. Filamentous *Xanthomonas oryzae* pv. *oryzae* were observed in bacterial aggregates associated with the protoxylem, scale bar: 10 µm. Post-processing was performed in GIMP (v2.10.24; pink: bacteria). **(C)**. Filamentous *Xanthomonas oryzae* pv. *oryzae* were associated with metaxylems, scale bar: 10 µm. Post-processing was performed in GIMP (v2.10.24; pink: bacteria). Related to Figures: Figure S6. Scanning electron microscopy of *Xanthomonas oryzae* pv. *oryzae* colonization *in planta* and vascular bundle structure in rice leaves.

**Figure 7.**
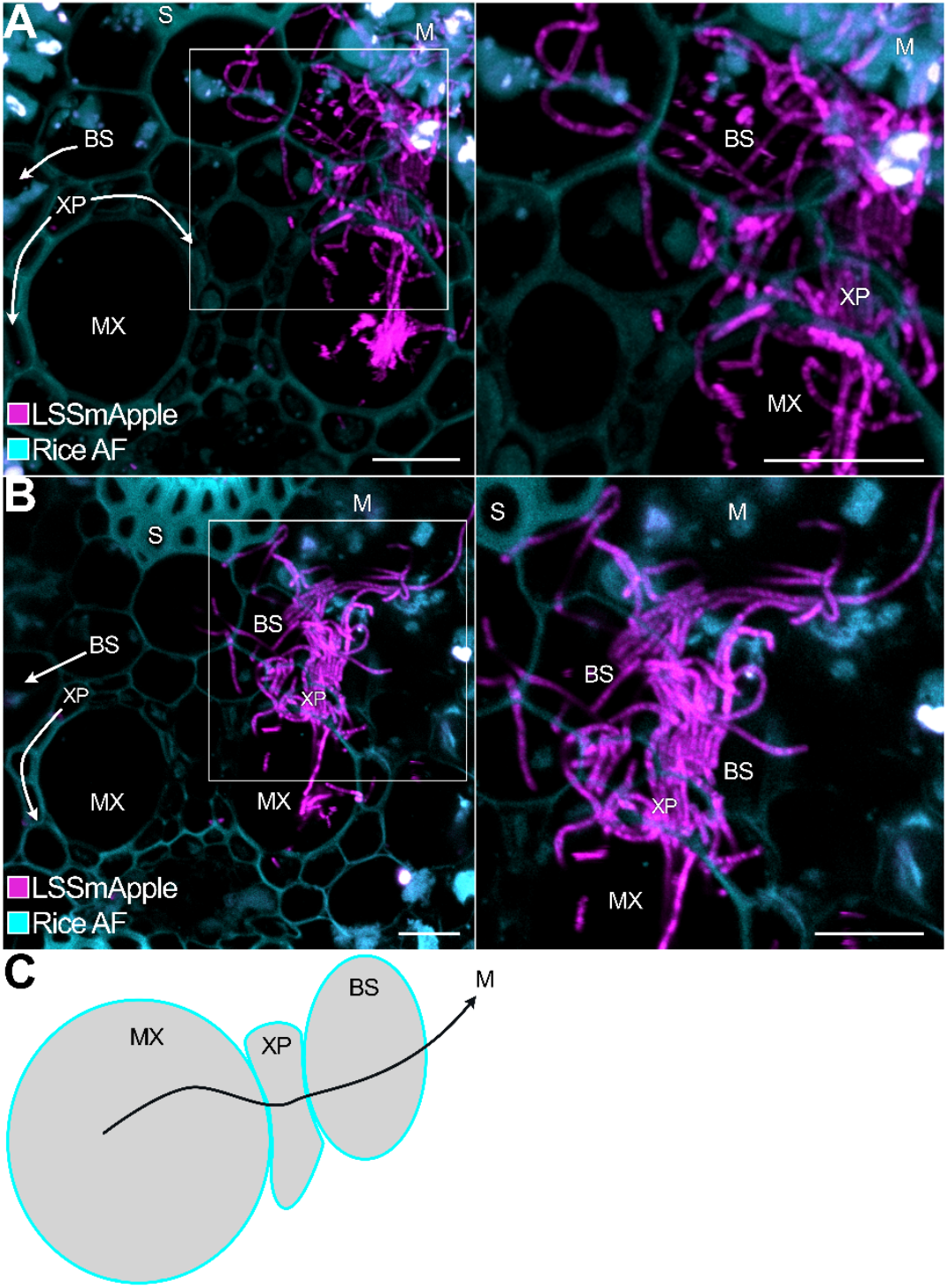
Filamentous *Xanthomonas oryzae* pv. *oryzae* traverse vascular bundle. Samples corresponding to an advanced stage of colonization (3-5 cm from the leaf tip at 6 dpi, Figure 4) were fixed and subjected CLSM microscopy. Experiment was performed three times independently with comparable results. **(A, B)**. Cross-section of *Xanthomonas oryzae* pv. *oryzae* during colonization, scale bar: 10 µm. Inlets indicated by white squares. PXO99^A^_LSSmApple_ (pink), rice autofluorescence (turquoise). **(C)**. Lateral migration route of Xoo from metaxylem vessels to xylem parenchyma and bundle sheath to mesophyll tissue. MX: metaxylem, XP: xylem parenchyma, BS: bundle sheath, M: mesophyll tissue, S: sclerenchyma. Related to Figures: Figure S4. Autofluorescence scans of rice leaves. Figure S7. Overview of rice leaf cross section. Figure S8. Colonization pattern of *Xanthomonas oryzae* pv. *oryzae* in cross sections of major veins visualized by CLSM. Figure S9. Colonization pattern of *Xanthomonas oryzae* pv. *oryzae* in cross sections of minor veins visualized by CLSM.

### Vascular bundle breach and lateral migration of filamentous Xoo at late stages

Notably, at late infection stages (6 dpi, 3-5 cm from infection site) SEM imaging revealed rod-shaped Xoo inside XP cells (Figure S6), likely a consequence of pit membrane rupture resulting from weakened cell walls due to xylan degradation by secreted Xoo xylanases, providing access to all the nutrients present in the dying XP (46). CLSM independently detected Xoo in XP (Figure 7). Notably, we found evidence for filamentous Xoo to progress from the xylem via the XP and bundle sheath even into the mesophyll (Figure 7, S7-9). Both rod shaped and filamentous cells were observed outside the xylem vessels. Filamentous cells traversed cell walls. How the cells achieved the penetration, and how, remains to be determined. We observed some variability in different leaves and in independent experiments, likely representing different stages of the progression of the infection (Figures S8, S9). In general, the depth of penetration was higher in primary veins as compared to secondary veins, consistent with the progression, in which primary veins were infected earlier compared to secondary veins as observed in the SWEET-GUS reporter assays (Figure 4). In some cases, the bacteria were still mainly present inside the xylem vessels; in other cases, penetrance into neighboring cells was observed (Figure S8A,B; S9A). In some cases, Xoo filaments accumulated highly in bundle sheath cells (S8 C,D, S9D), and in other cases, the filaments entered the mesophyll (mesophyll cell identifiable by high chlorophyll autofluorescence)(Figure S8C,D; S9D). We surmise that the differences are due to differences in the stage of infection, and that they represent representative illustration of the progression from xylem vessels to XP to bundle sheath and ultimately the mesophyll. Thus, rather than infecting rice leaves in a single static morphology, Xoo undergoes dynamic shape transitions *in planta*, processes with likely central importance for virulence.

## Discussion

Using a combination of wide filed, SEM and CLSM imaging of Xoo labelled with the large Stokes shift fluorescent protein LSSmApple, we found that Xoo can differentiate to produce morphologically different forms, from rod-shaped bacilli to non-septated filaments to septated filaments both *in vitro* and during the infection of rice. Filamentation can be triggered *in vitro* by extreme conditions such as high salinity or high density, however the filamentation is reversible. Colony-forming ability was shown to differ between bacilli and filaments within the bacterial population. These reversible transitions may have a varieties of benefits during the infection process, e.g. by increasing the surface that can attach to the xylem walls to withstand the xylem sap flow to penetration of host cell layers to spread beyond the vascular bundle, demonstrating that Xoo is a xylem-spreading pathogen that at later stages of the infection can become necrotrophic, and thus can be defined as a hemi-biotroph. Not surprisingly, Xoo is therefore in some aspects similar to the closely related non-vascular *Xanthomonas oryzae pv. oryzicola*, which differs from Xoo not having CsbA, a cellobiohydrolase gene, as a key determinant of the difference between vascular and non-vascular life styles (47). Vascular and non-vascular pathogens of *Brassicaceae, Xanthomonas campestris* pv. *campestris* (Xcc) and pv. *raphani* differ in their CRISPR/Cas repertoires. Loss of CRISPR/Cas increased genome plasticity and enabled the accumulate virulence factors (e.g. XopN) that enabled niche adaption of Xcc to vasculature (48). Notably, these findings may be of relevance for other xylem pathogens and possibly beyond plant diseases.

### Xoo is not xylem-limited and displays phenotypic heterogeneity during vascular bundle breach

*Xanthomonas* species encompass vascular and non-vascular pathogens. Non-vascular pathogens e.g. *Xanthomonas oryzae* pv. *oryzicola* invade plants through stomata and colonize mesophyll tissue. Vascular pathogens e.g. *Xanthomonas campestris* pv. *campestris* (Xcc) and Xoo are able to colonize xylem vasculature despite its scarce nutrient availability and the rapidly flowing xylem stream. At stomata, immune hubs like BAK1/BKK1 interact with pattern recognition receptors for specific pathogen cues e.g. flagellin and mediate stomata closure. At hydathodes, defense layers of plant immunity appear to be restricted to immune hubs and lack specific pattern recognition receptors (49,50). It has been hypothesized that vascular pathogens hijack guttation droplets at hydathodes to be reabsorbed into the epithem when xylem pressure decreases under low water availability and low humidity. While initial plant invasion into the epithem appears to be less demanding for vascular pathogens because of the limited plant defense layers at hydathodes, Type II Secretion System (T2SS) secreted cell wall degrading enzymes (CWDEs) are essential for the degradation of xylem tracheid apices and progression into xylem vasculature from the epithem (49). To our knowledge, the ability of the vascular-spreading Xoo to not only gain entry into the xylem vasculature but also exit the xylem vessels had not been reported before. Our observations indicate that Xoo is not restricted to basipetal migration along xylem vasculature (Figure 7). For colony expansion beyond xylem vasculature, Xoo is required to surpass pit elements of the xylem vessel. Xylem pits are composed of a pliant pit membrane (pectin and hemicelluloses e.g. heteroxylan) in contact with the plasma membrane of adjacent xylem parenchyma cells (51,52). Plausibly, the CWDEs essential for xylem vasculature entry after epithem colonization can also degrade xylem pits and enable pathogen spread to xylem parenchyma cells (47,53). Transcriptomic studies of Xoo reported the upregulation of xylanases *in planta* (46,54), which enable Xoo to degrade pits composed of heteroxylan (52) to xylose and provide a passageway to adjacent XP cells. In addition, we observed that pleomorphic adaption into filamentous phenotypes appeared crucial for migration outside the vascular bundle: filaments spread from xylem vessels to XP and bundle sheath cells to mesophyll tissue (Figure 7) Individuals of the same species can exhibit different behaviors of so-called phenotypic heterogeneity within one colony, benefitting the population rather than the individual (bet-hedging strategy). Division of labor, achieved by heterogeneity has been reported for plant-pathogens (22,39,55). Filamentous Xoo were observed at advanced infection stages, where bacterial cell density probably culminated within the xylem vessel, which could be attributed to quorum-sensing signals, as seen in *X. fastidiosa* where cell density and exposure to quorum-sensing signals triggered filamentation (22). Pleomorphic bacteria could possibly perform distinct roles. For instance, filaments could bridge the gap between separated subpopulations within xylem vessels for basipetal migration and outside the vascular bundle, similar to observations of *X. fastidiosa ex planta* (22). In *X. fastidiosa*, filamentous cells act as anchors of biofilm due to their irreversible attachment to surfaces and an enhanced production of extracellular polymeric substances (34). Due to their increased surface, an increased amount of afimbrial adhesins could mediate adherence to adjacent colonization sites. In contrast to the potential anchoring role of filaments at high cell density, the increased length of filamentous bacteria could fit significantly more pili compared to rod-shaped bacilli and fulfill a dispersal role within the bacterial population. Filamentous bacteria have been associated with rapid dispersal via swarming (42,56–59). Since we observed the ability of Xoo filaments to found new pleomorphic colonies by differentiation and division (Figure 3), filamentous dispersal could lead to efficient and rapid colony expansion. Whether Xoo filaments fulfill specific tasks within the bacterial population associated to swarming and dispersal or non-motile anchoring remains to be investigated. Filaments of other bacterial species have been associated with both, non-motility (39) or dispersal (22,41,56–59).

Although the mechanisms and intracellular signaling pathways underlying filamentation remain poorly defined, diverse cues ultimately converge to regulate the cell division machinery. For instance, the SOS response and the UDP-glucose pathway induce inhibitors of FtsZ ring assembly and lead to lateral growth but inhibited cell division (60–64). Filaments of *Bordetella antropi* were reported to be induced under high UDP-glucose conditions enabling cell-to-cell dispersal (40). Hence, a next step is to explore the effect of the nutrient status on Xoo filamentation. During primary infection, Xoo redirects the sucrose flow via induction of *SWEETs* to the site of infection at pits. One may hypothesize that at later stages of infection, XP sucrose reservoirs may have been exploited. While nutrient excess will boost Xoo proliferation, Xoo cells within the same microcolony will likely compete for nutrients at high cell density. Effects of clustered growth of filaments has been discussed as primarily negative due to impaired motility and increased competition for resources (65). In other bacteria, low nutrient levels (16,17,66), as well as conditions with high nutrient levels have been described to induce filamentation (18,39).

### Infection front predominated by rod-shaped *Xoo* at xylem vessel pits

In the xylem vessels, Xoo faces low microbial competition (51,67) but needs to withstand and migrate against the bulk laminar flow of the xylem sap. Bacterial attachment to xylem vessel walls is mediated by adhesins (68,69). Bacterial migration within the xylem vessel occurs bidirectionally by type IV twitching and flagellar swimming (68,70). We found that at the infection front, Xoo frequently localized in xylem pits, and many cell aligned perpendicular to xylem vessel pits (Figure 5). Likely, pit membranes, which are composed of hemicellulose are an easier target for Xoo T3SS injection compared to the stiff and highly suberized secondary cell walls of the xylem vessel walls (52). 45 % of Xoo cells appeared to be orientated perpendicular to pit membranes, indicating that adherence and possibly T3SS secretion are polarized (Figure 5). Upon injection of TALe, *SWEET* induction redirects the sucrose flux to the nutrient scarce xylem sap. When the plant cannot contain Xoo infection by deposition of tyloses, thickening of xylem vessel walls or other defense responses (4,71), Xoo migration will progress.

### Route of Xoo infection detected with translational SWEET-GUS reporter lines

To date, the colonization route of Xoo during infection was deduced from the appearance of the chlorotic (pale yellow) and necrotrophic (grey) lesions after leaf clip infection (72). Our analysis here relied on the pathogen’s requirement for *SWEET* gene induction and resolved colonization patterns more sensitively than lesion lengths. Translational SWEET11a-GUS reporter lines enabled accurate spatiotemporal resolution of bacterial distribution, revealing a basipetal infection pattern from main veins through transverse veins to minor veins. Notably the infection process was found to be non-contiguous, but to occur via formation of new colonies that migrated basipetally. Only strains carrying the SWEET11a-inducing TAL effector PthXo1 induced diX Indigo accumulation, confirming the effector dependency of the Xoo-rice pathosystem (6,73). The migration of Xoo within rice vasculature corresponds to leaf vein architecture and likely reflects a strategy for systemic spread and nutrient acquisition. Mapping of the Xoo infection route helped us to examine specific phases of colonization using CLSM and SEM.

### Conditional filamentation of Xoo lead to different viability of filaments *in vitro*

On agar plates, Xoo colonies developed a zonal architecture with three concentric regions. Each zone displayed different optical properties and distinct mixtures of rod-shaped bacilli and filamentous cells (Figure 1). The colony center contained mostly rod-shaped cells, while the periphery was enriched in filamentous forms. This spatial organization likely reflects gradients in nutrients, oxygen, and cell signaling molecules. Filaments were also observed for Xoo exposed to high salinity or high OD_600_ in liquid cultures (Figure 2). Our findings are consistent with bacterial pathogen responses to environmental conditions, where filamentation often serves as an conditional phenotypic change to enhance resilience (19), compared to obligate filamentous species (74–76). When populations were separated by cell length and growth history, filamentous cells from high salinity cultures produced the highest number of CFU, indicative of high active reproduction (Figure 3). By contrast, filamentous cells from high-density cultures formed fewer colonies. Our data imply that depending on the status of the originator population, filamentous cells can share the same phenotypic features but differ in their viability. Filamentation in Xoo appears to represent a conditional response of bacteria to environmental conditions. The *in vitro* system may allow us in the next step to identify differences in gene expression in bacilli and filamentous stages, e.g. by RNAseq, as a basis for generating mutants to explore the mechanism of reversible filamentation and their role in host colonization.

### Xoo filaments can revert into pleomorphic colonies

To gain insights into the developmental program of filamentation, filamentous cells were observed by time lapse over a 24-hour period in a live imaging set-up. Filamentous cells elongated after transferal to imaging chamber, subsequently forming septa at irregular positions and then divided asymmetrically (Figure 3). The divisions generated a mixed population containing rod-shaped, intermediate, and filamentous cells, as described for other bacteria (21,77,78). The reversible transition of filamentous to pleomorphic cell morphology indicates that filamentation represents a transient stage rather than a terminal state. In contrast to the unipolar growth observed for *Corynebacteriu matruchotii* and *Leptrothis cholodnii* (16,79), bipolar peptidoglycan synthesis appeared to occur in Xoo. In the context of host colonization, reversible filamentation could allow the mobilization of cells for further colonization in the basipetal direction in leaves and the foundation of new colonies, as observed in the SWEET-GUS-reporter system population expansion. Mutants defective in the different steps of this cycle in conjunction with the reporter systems developed here may help to gain insights into the exact process.

### Conclusion

The patterns of Xoo morphotypes are indicative of coordinated bacterial behavior that enhances nutrient acquisition and adhesion, potentially through subpopulation-specific traits governed by cell length within the population. Given that Xoo must overcome physical barriers within the host, including rapid xylem flow velocities and immune responses, morphological plasticity may provide distinct ecological and survival advantages. For example, TAL effector delivery likely occurs preferentially at xylem vessel pits where rod-shaped cells adherence perpendicularly, while filamentous cells appear to dominate during lateral spread between vascular bundles and systemic infection to mesophyll tissue. In conclusion, our study provides compelling evidence for the adaptive significance of morphological plasticity in Xoo, particularly filamentation, in response to environmental cues and during host colonization. This morphological flexibility likely represents a critical component of Xoo’s infection strategy, enabling it to navigate and persist within the physically challenging and nutritionally constrained environment of rice xylem vessels. Understanding the regulatory mechanisms underlying morphological transitions and their impacts on bacterial physiology and virulence may inform future strategies for managing bacterial blight in rice, but also other bacterial diseases.

## Data availability

Raw and meta data are available at: https://doi.org/10.60534/8d4s9-ywr51

## Acknowledgements

We thank Joon Seob Eom for initial experiments on infection progression in rice leaves as monitored using a stably transformed translational SWEET11a-GUS line. We also thank Sebastian Hänsch and Steffen Köhler from the Center for Advanced Imaging (CAi) for training, continuous advice and support with both confocal and SEM imaging and data analysis. We thank Dominik Brilhaus (CEPLAS) for training and advice on ARC generation, publication of raw and meta data and Tom Boissonnet (CAi) for microscopy data management. This work was supported by grants from Deutsche Forschungsgemeinschaft (DFG, German Research Foundation) - Collaborative Research Center SFB1535, project ID 458090666/CRC1535/1; Deutsche Forschungsgemeinschaft (DFG, German Research Foundation) under Germany’s Excellence Strategy – EXC-2048/1 – project ID 390686111 (CEPLAS2), and the Alexander von Humboldt Professorship to WF. LR was supported by a stipend from the Heinrich Böll Stiftung and CEPLAS.

## Declaration of interests

The authors declare no competing interests.

## Author contributions

LR and WBF conceived the study. LR, ZYM and AR performed experiments. VSL and EL supervised the team. MB supported LR for SEM on rice leaves. LR, ZYM and WBF performed analyses. LR and WBF wrote the manuscript.

## STAR★Methods

### Key resources table

| REAGENT or RESOURCE | SOURCE | IDENTIFIER |
| --- | --- | --- |
| **Bacterial strains** |  |  |
| PXO99<sup>A</sup> | Lab of Boris Szurek, IRD | PXO99<sup>A</sup> |
| PXO99<sup>A</sup><sub>LSSmApple</sub> | This paper | PXO99<sup>A</sup><sub>LSSmApple</sub> |
| ME2 | Lab of Boris Szurek, IRD | ME2 |
| **Experimental models: organisms/strains** |  |  |
| *Oryza sativa* L. subsp. *japonica* cv. Kitaake | Eom et al. 2019 | Wild type Kitaake |
| *Oryza sativa* L. subsp. *japonica* cv. Kitaake, pSWEET11a:SWEET11a-GUSplus T4, transformation event 10 | Eom et al. 2019 | SWEET11a-GUS, #10 |
| *Oryza sativa* L. subsp. *japonica* cv. Kitaake, pSWEET11a:SWEET11a-GUSplus T4, transformation event 8 | Eom et al. 2019 | SWEET11a-GUS, #8 |
| **Recombinant DNA** |  |  |
| pNEO:GFP | Han et al. 2008 |  |
| LSSmApple | Ejike et al. 2023 |  |
| **Software** |  |  |
| R v4.20 | R core team | www.r-project.org/ |
| Affinity Designer | Serif | https://affinity.serif.com |
| GIMP | GIMP's Team | www.gimp.org/ |
| OMERO | Center for advanced Imaging, HHU | https://cai.hhu.de/omero |
| **Other** |  |  |
| Zeiss LSM 880 Airyscan | Center for advanced Imaging, HHU |  |
| Leica TCS SP8 | Center for advanced Imaging, HHU |  |
| Zeiss Supra 55 VP | Center for advanced Imaging, HHU |  |
| Nikon Ti Eclipse PF | Center for advanced Imaging, HHU |  |

## Experimental model and subject details

### *Xanthomonas oryzae* pv. *oryzae* culture conditions

Bacterial pre-cultures were obtained from single colonies derived from glycerol stocks streaked on NBSA plates, incubated for two days at 28°C in dark conditions (Table S1). 5 µL of bacterial pre-culture at OD_600_ 0.4 were transferred into 300 mL Erlenmeyer flasks, filled to a final volume of 50 mL NBS (1 g/L yeast extract, 3 g/L beef extract, 5 g/L peptone, 10 g/L sucrose, pH 7, 100 µg/mL kanamycin in distilled H_2_O) or 500 mM NaCl in NBS. After an initial 16 hours of incubation (150 rpm, 28 °C, dark conditions), growth of cultures was monitored hourly by optical density at OD_600_. At OD_600_ 0.4 for control and 500 mM NaCl conditions and OD_600_ 3 for high cell density conditions, bacteria were collected in ice-cold falcon tubes and centrifuged (1500 g, 4°C) for 15 min.

### Plant cultivation

Rice seeds were de-husked and sterilized on a shaker at 180 rpm in 15 ml Falcon tubes filled with 75 % Ethanol and 50 % Klorix® for 2 and 5 min, respectively (Table S2). The seeds were dried on sterile filter papers and transferred onto Magenta™GA-7 boxes containing ½ salt strength MS media (2.2 g Murashige Skoog Medium, 10 g sucrose, and 8 g Phytagel per liter, pH 5.8) with a serological tweezer under aseptic conditions. For SWEET-GUS reporter lines seeds from transformation event 8 and 10 (6), medium was supplemented with 50 mg/L hygromycin B (Table S2). Seedlings were grown in a long day light regime (16 h day/8 h night) with a photosynthetic photon flux density (PPFD) of 200 µmol m^-2^ s^-1^ (PPFD-blue 40, PPFD-green 80 and PPFD-red 70 µmol m^-2^ s^-1^), at 27°C and 80 % humidity in CLF PlantClimatics chambers (model: CU41L5).

## Method details

### Histochemical GUS assay

The newly emerged leaf of translational GUS reporter lines at the 4^th^ leaf stage was infected with PXO99^A^ and ME2 inoculum after Kaufmann et al. (1973). Samples for histology were collected in ice-cold 90 % acetone and vacuum infiltrated for 10 min and incubated for 30 min. Subsequently, the fixing solution was replaced by GUS washing buffer followed by staining buffer and were infiltrated for 10 min each (Table S3). After incubation at 37°C, the enzymatic GUS reaction was stopped by replacement of GUS staining buffer with 75°C ethanol. Samples were cleared with a series of 75 %, 80 % and 90 % ethanol (6). Final clearing was performed with 15 % chloral hydrate. Images were taken with the AxioZoom.V16 (Zeiss) using stitching in bright field mode.

### Generation of fluorescent *Xanthomonas oryzae* pv. *oryzae*

The large Stokes shift protein LSSmApple was integrated into the expression plasmid pNEO via In-Fusion Assembly Mix (Takara) in frame with the neomycin promoter (80,81). After successful assembly of the pNEO:LSSmApple, the electrocompetent Xoo strain PXO99^A^ was transformed with 500 ng of plasmid DNA for 5 seconds at 2.5 kV and 200 Ω in a 0.1 cm cuvette. Cells were recovered in Recovery Medium for Expression (Sigma, CMR001-8X12ML) for 2 hours shaken at 28 °C and plated on NBSA (1 g/L yest extract, 3 g/L beef extract, 5 g/L peptone, 10 g/L sucrose, pH 7, 1.5% w/v agar, 100 µg/mL kanamycin in distilled H_2_O) and stored at 28 °C in dark conditions for four days. Kanamycin tolerant transformants were selected for fluorescence using a fluorescent Stereo Zoom Microscope (AxioZoom.V16, Zeiss) and, LSSmApple filter (excitation 488/10, emisson605/50, beamsplitter zt514). Fluorescent transformants were tested for virulence using the leaf clip infection protocol after Kaufmann et al. (1973).

### Autofluorescence scan of rice leaves

Emission scans were acquired on a Leica TCS SP8 confocal microscope using a 40× water-immersion objective. Excitation was set to 405, 488, 514, and 561 nm, with notch filters applied when available to reduce laser bleed-through. Emission was collected from excitation + 20 nm to 640 nm, or up to 740 nm when chloroplast autofluorescence was included, in 10 nm steps using the PMT spectral detector. The pinhole was set to 2 Airy unit and the scan speed to 400 Hz. Detector gain and laser power were adjusted to avoid saturation across the scan. Regions of interest were selected on both the adaxial and abaxial sides of the 4th leaf. Three biological replicates were analyzed.

### Preparation of infected rice samples, microscopy and image processing

Infected leaf samples were collected and submerged in fixation solution (freshly prepared 4 % PFA, 0.1 % Triton X-100 in PBS) shaken at 80 rpm, 4°C for 16 hours. Samples were washed three times in 0.1 Triton X-100 PBS as well as PBS for 10 min and chopped roughly into pieces with a razor blade. Samples were collected in a 16 well dish filled with PBS and cut into thin cross or longitudinal sections. CSLM of rice leaf sections was performed with the Zeiss LSM880 with a Plan-Apochromat 40x/1.2 objective. For selection of the infection front, infected rice leaves were observed with a fluorescent Stereo Zoom Microscope (AxioZoom.V16, Zeiss) and, LSSmApple filter (excitation 488/10, emisson605/50, beamsplitter zt514) and 63 HE mRFP filter (excitation 565/30, emission: 620/60, beamspliter FT585) for rice autofluorescence. Samples were taken 0.5 cm from front of bacteria (Figure S6). exposed to a series of ethanol concentrations after fixation and washing, from 10 % ascending by 5 % up to 99.9 % for 15 min each. After critical point drying, samples were coated with gold using an Agar Sputter Coater and imaged with a Zeiss SUPRA 55 VP SEM (EHT = 5 kV). Post-coloration of SEM images was performed in GIMP (v2.10.24).

### Bacterial viability assay

After an initial 16 hours of incubation (150 rpm, 28 °C, dark conditions), growth performance of cultures was monitored hourly by OD_600_. At OD_600_ 0.4 for control and 500 mM NaCl conditions and OD_600_ 3 for high cell density conditions, bacteria were collected in ice-cold falcon tubes and centrifuged (1500 g, 4°C) for 15 min. To test the colony forming ability of Xoo cells of different length, bacterial pellets were diluted in 150 mL of ice-cold NB (1 g/L yest extract, 3 g/L beef extract, 5 g/L peptone, pH 7, 100 µg/mL kanamycin in distilled H_2_O) and filtered through 5 and 10 µm pluriStrainer into falcon tubes on ice, respectively. Bacterial flow through was centrifuged (15 min, 1500 g, 4°C) and diluted in NBS to OD_600_ 0.1. For the CFU assay, subpopulations of bacteria at OD_600_ 0.1 were diluted by 10^6^ in NBS. 30 µL of diluted bacterial suspensions were evenly spread with sterile metal beads on NBSA plates supplemented with 100 µg/mL Kanamycin. CFU were quantified after three days of incubation at 28°C in dark conditions.

### Microscopy of *Xanthomonas oryzae* pv. *oryzae*

For long-term life imaging of Xoo, 10 mL of bacterial culture was centrifuged for 15 min at 1500 g, 4 °C. The bacterial pellet was eluted gently with 1 mL NBS media. 5 µL of bacterial suspension was transferred to an agar-based aerated imaging chamber sealed with silicon-based grease (Fig S4). A z-stack image was acquired every 30 min for 24 hours with the Nikon Ti Eclipse PFS. For high resolution images of Xoo cells after filamentation, samples were taken from 0.5, 2, 4, 8, 10 and 16 hours after transferal to imaging chamber with CLSM (excitation 488 nm, emission 605-615 nm; Plan-Apochromat 63x/1.4 objective). Analysis was performed in Omero (v5.28.0) and ImageJ (v1.52).

## Notes

https://doi.org/10.60534/8d4s9-ywr51

### Competing Interest Statement

The authors have declared no competing interest.

