## Supplementary material for "Differentiation of *Xanthomonas oryzae* pv. *oryzae in vitro* and during rice leaf infection": Figure S1

Figure S1. HSB color space segmentation of bacterial colonies.
Figure S2. Assembly of an agar-based aerated imaging chamber.
Figure S3. Time-lapse imaging of filamentous *Xanthomonas oryzae* pv. *oryzae*.
Figure S4. Autofluorescence scans of rice leaves.
Figure S5. Selection of infection front for SEM.
Figure S6. Scanning electron microscopy of *Xanthomonas oryzae* pv. *oryzae* colonization in planta and vascular bundle structure in rice leaves.
Figure S7. Overview of rice leaf cross section.
Figure S8. Colonization pattern of *Xanthomonas oryzae* pv. *oryzae* in cross sections of major veins visualized by CLSM
Figure S9. Colonization pattern of *Xanthomonas oryzae* pv. *oryzae* in cross sections of minor veins visualized by CLSM.

Tabel S1. Xoo strains used in this study
Table S2 Rice lines used in this study.
Table S3. Recipes for solutions for GUS histochemistry

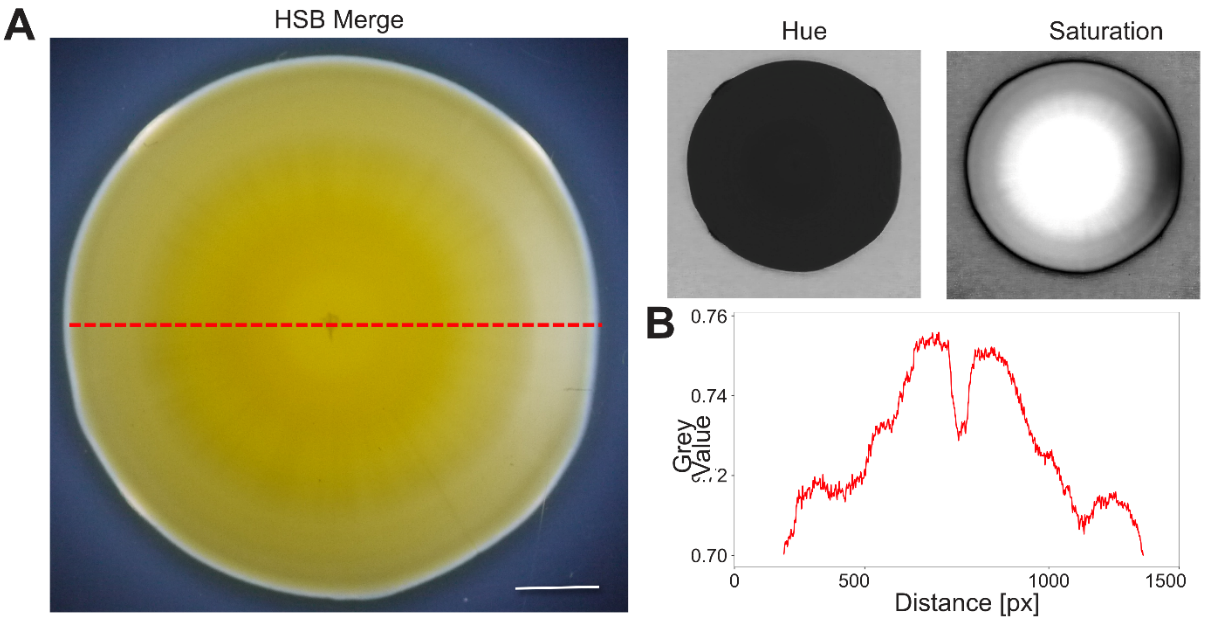


**Figure S1. HSB color space segmentation of bacterial colonies.**
**(A).** Colonies were analyzed in the hue-saturation-brightness (HSB) color space to enable pixel-level discrimination of structural and optical features across the colony surface. **(B).** Segmentation was based on the grayscale value of pixels analyzed for pixels on the red-line in the HSB merge image.

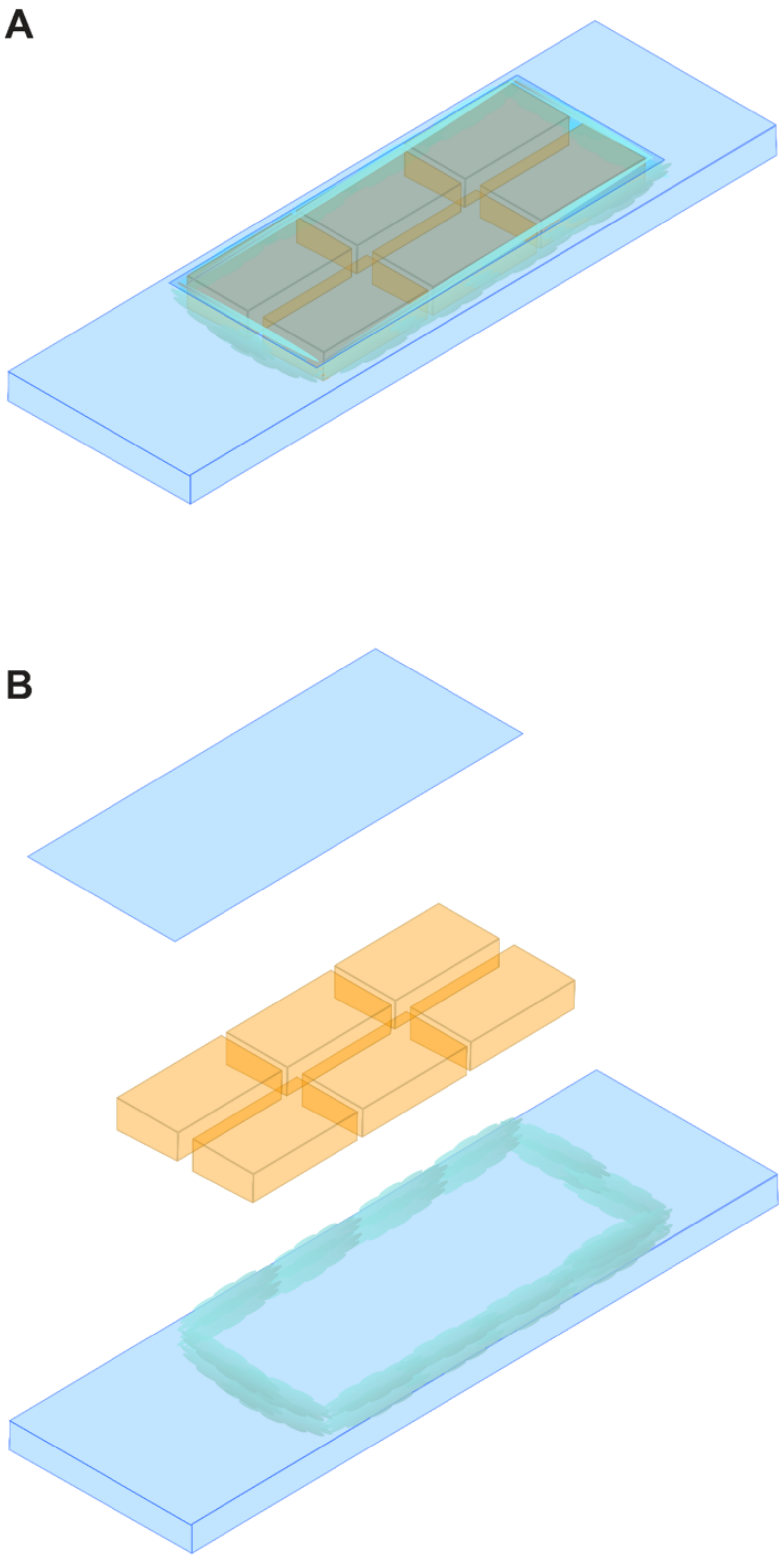


**Figure S2. Assembly of an agar-based aerated imaging chamber.** The chamber was constructed by placing six thin, evenly poured NBSA media blocks onto a glass slide to in 0.5 cm distance to maintain aeration. Imaging dish and cover glass was surrounded with a rim of silicon grease to prevent desiccation. Long-term live-cell imaging was performed with Zeiss Immersol W immersion liquid.

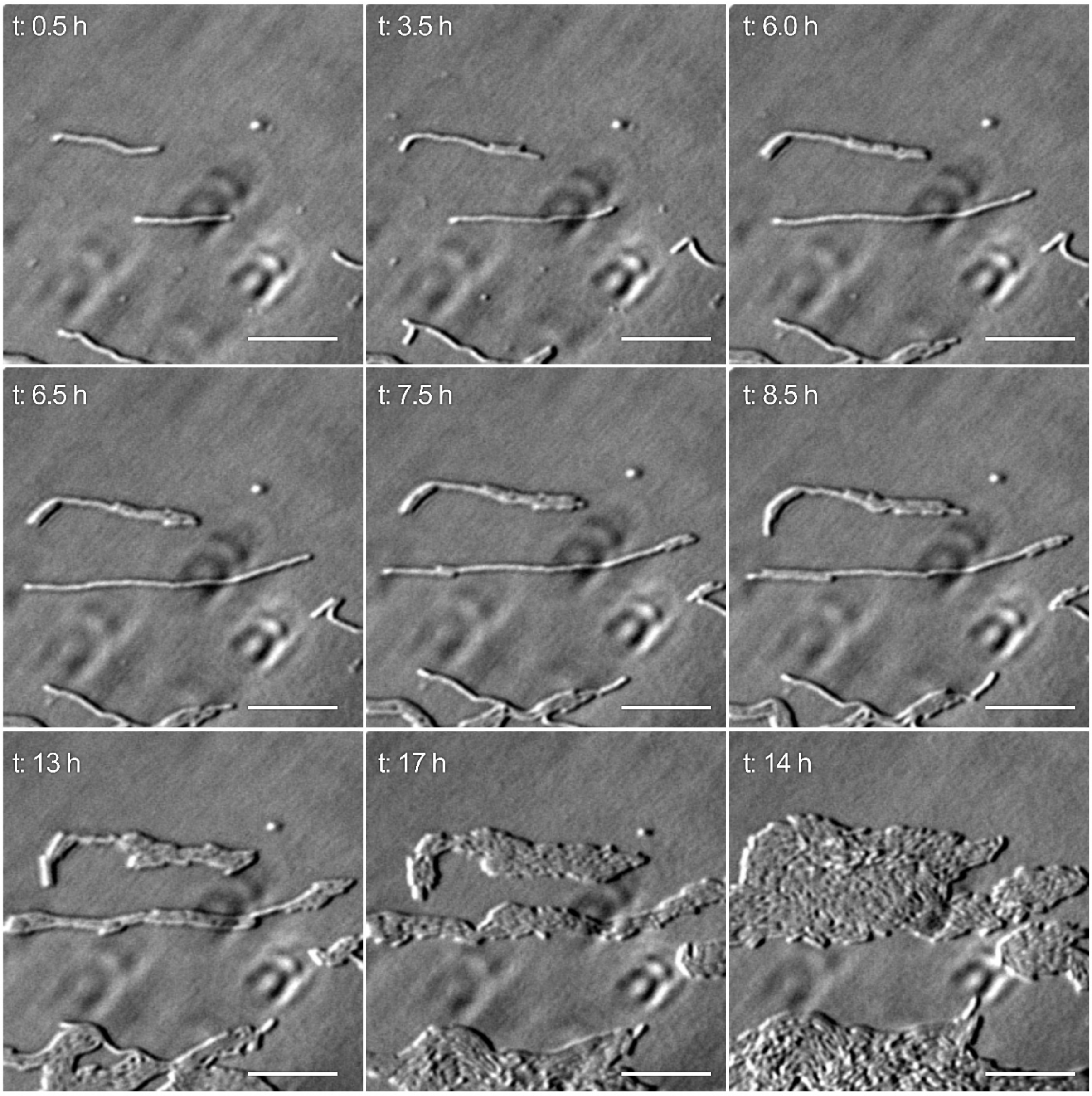


**Figure S3. Time-lapse imaging of filamentous *Xanthomonas oryzae* pv. *oryzae*.** Filamentous cells were separated by filtration from cultures grown in high salinity ($OD_{600}$ 0.4, 500 mM NaCl) and transferred onto NBSA medium within the aerated imaging chamber (Figure S4). Individual filaments were monitored for 24 h by bright-field microscopy (Nikon Ti Eclipse PFS). Representative images at selected time points are shown; Z-stacks were acquired every 30 min for 24 h. Scale bar: 10 µm.

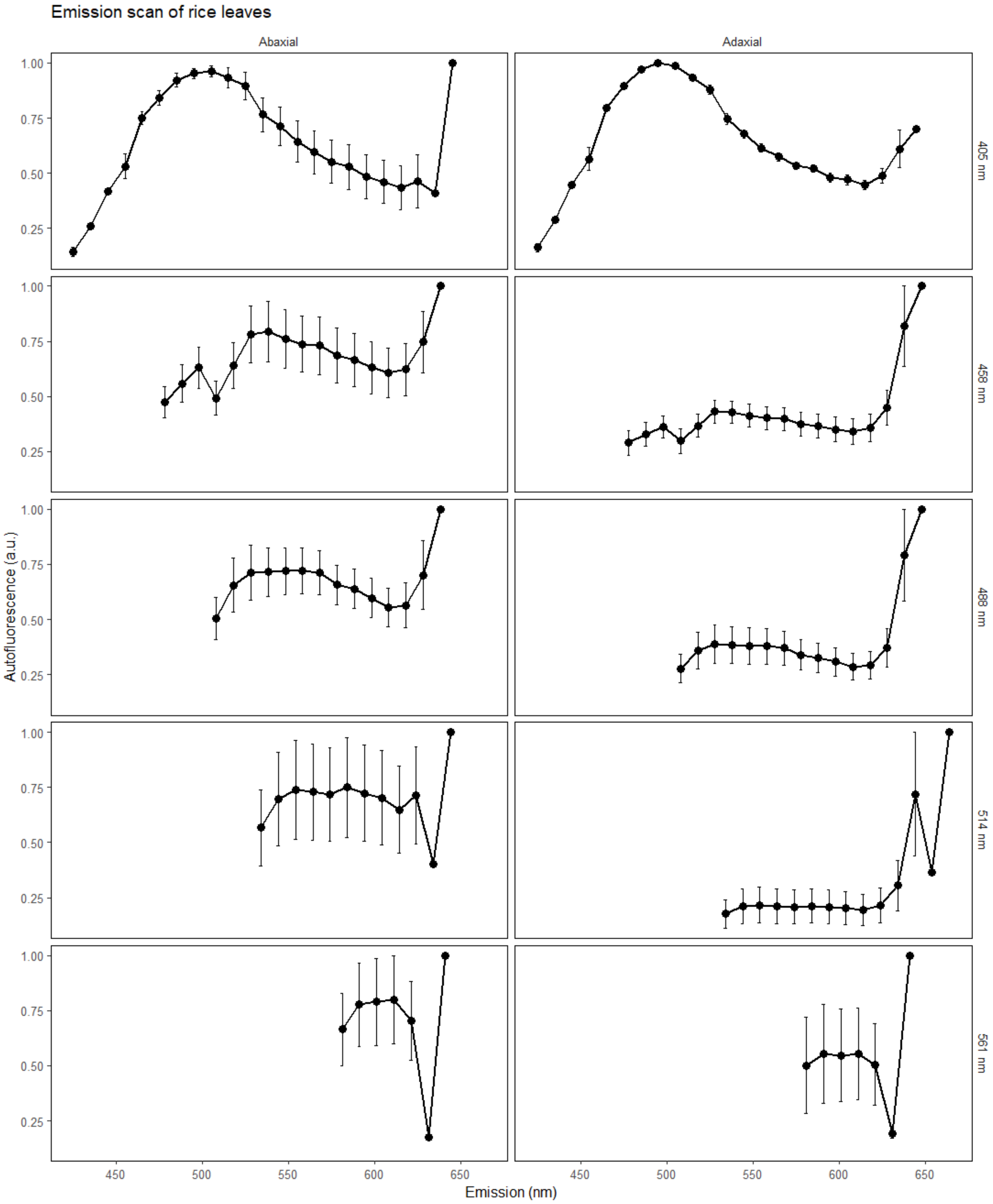


**Figure S4. Autofluorescence scans of rice leaves.**
Emission spectra were acquired on a Leica TCS SP8 using a 40× water-immersion objective with excitation at 405, 488, 514, and 561 nm. Emission was collected in 10 nm steps from excitation +20 nm up to 640 nm (peak at 740 nm: chloroplast autofluorescence). Regions of interest were recorded on adaxial and abaxial surfaces of the fourth leaf, with three biological replicates analyzed.

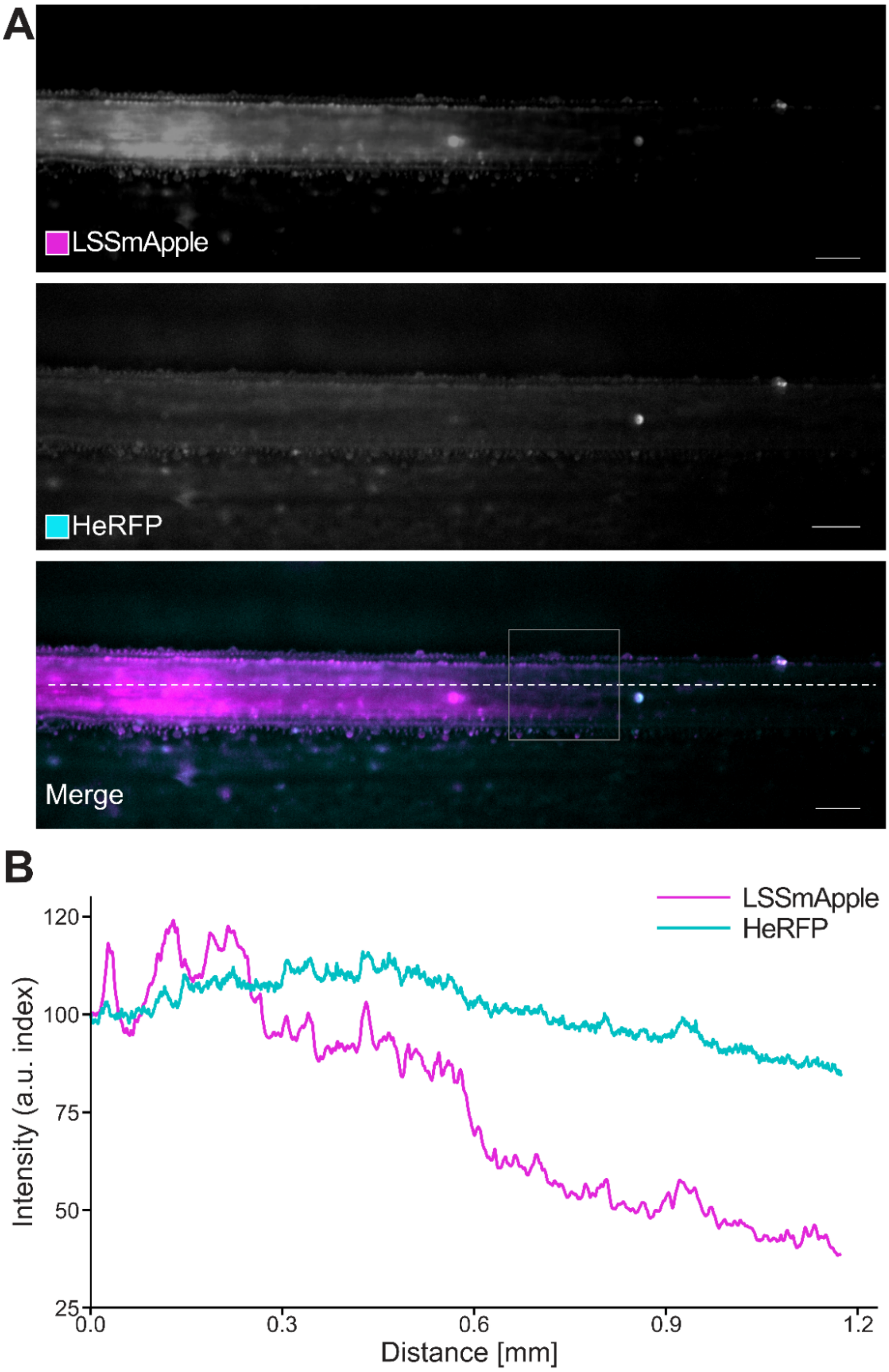


**Figure S5. Identification of the infection front for SEM with PXO99$^{A}_{LSSmApple}$.** For scanning electron images of the infection front, regions spanning at ~0.5 cm from the visible infection front observed by PXO99$^{A}_{LSSmApple}$ fluorescence signal were selected. (A) Images of rice xylem colonized by PXO99$^{A}_{LSSmApple}$. Selected region for processed for SEM analyzes indicated by square (B) Fluorescence intensity profile of LSSmApple compared to rice autofluorescence detected with 63 HE mRFP filter (indicated by dashed line in A).

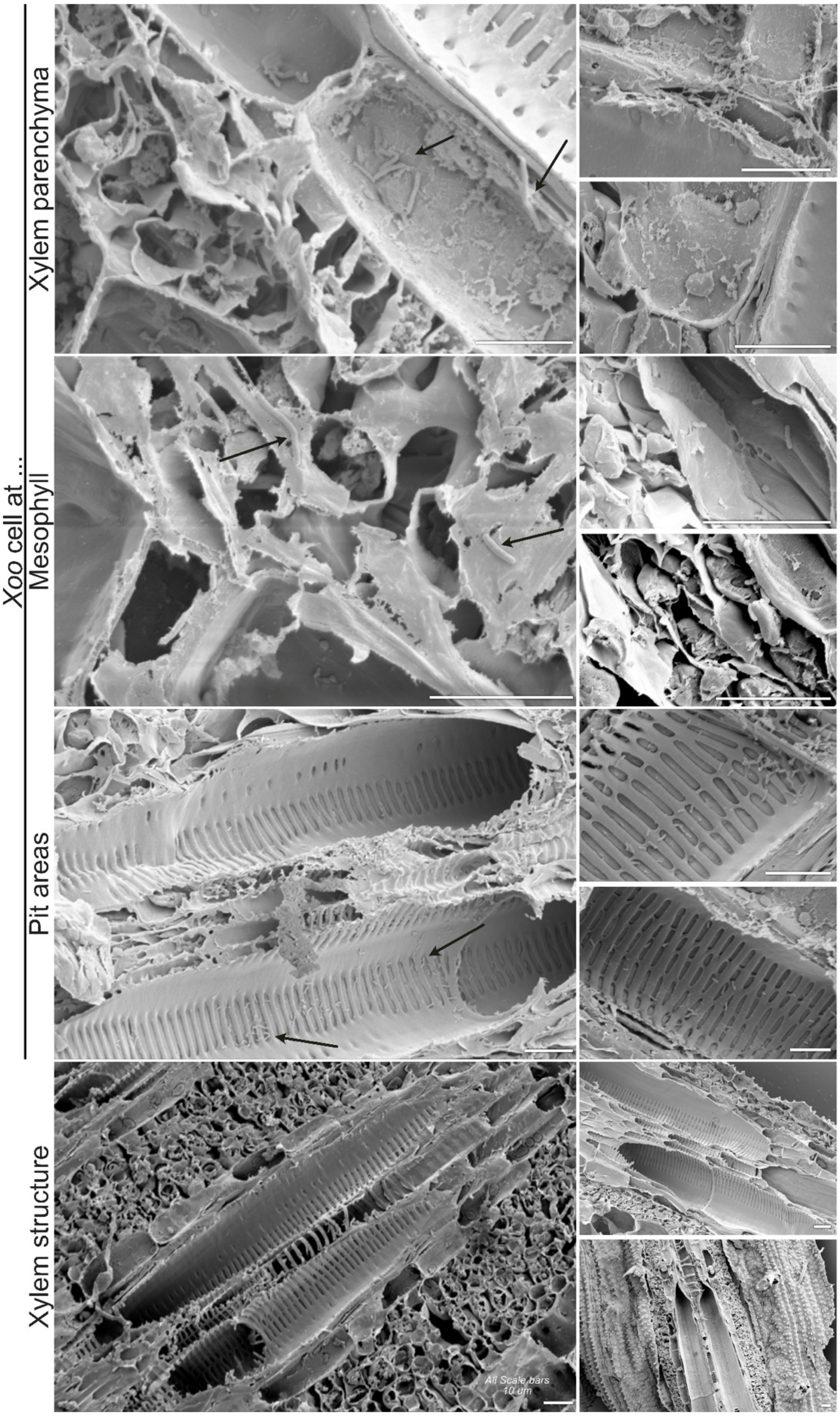


**Figure S6. Scanning electron microscopy of *Xanthomonas oryzae* pv. *oryzae* colonization *in planta* and vascular bundle structure in rice leaves.** Representative SEM images showing Xoo localization and xylem ultrastructure in infected rice leaves. Xoo cells (black arrows) within XP (first panel). Xoo cells detected in mesophyll tissue (second panel). Xoo cells at xylem pits in metaxylem vessels (Third panel). Detailed view of overall vascular bundle architecture in rice, highlighting metaxylem vessels, tracheids and xylem parenchyma cells of the xylem (fourth panel). Scale bars: 10 µm.

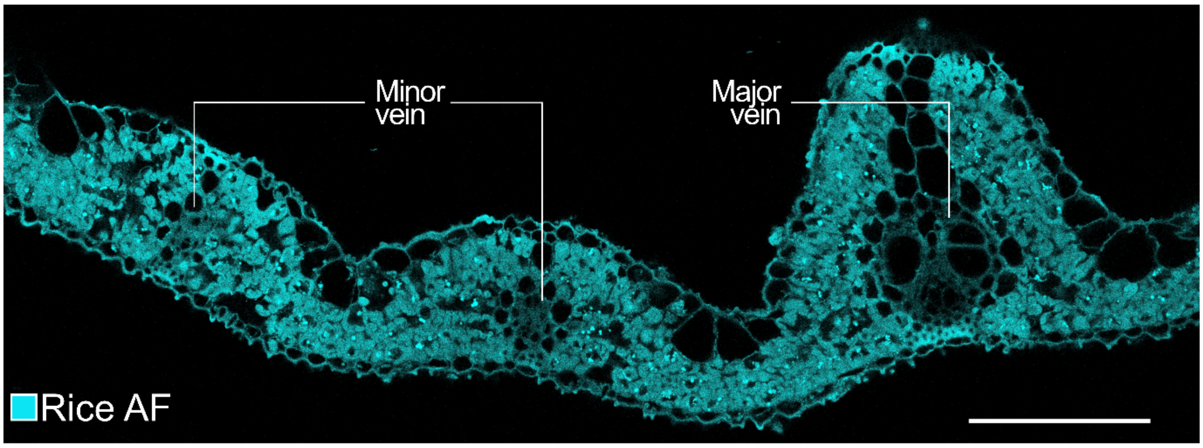


**Figure S7. Overview of rice leaf cross section.**
Cross section of rice leaf. Shown here: two secondary veins and a primary vein. Rice autofluorescence at 488 nm excitation shown in turquoise. Scale bar: 100 µm.

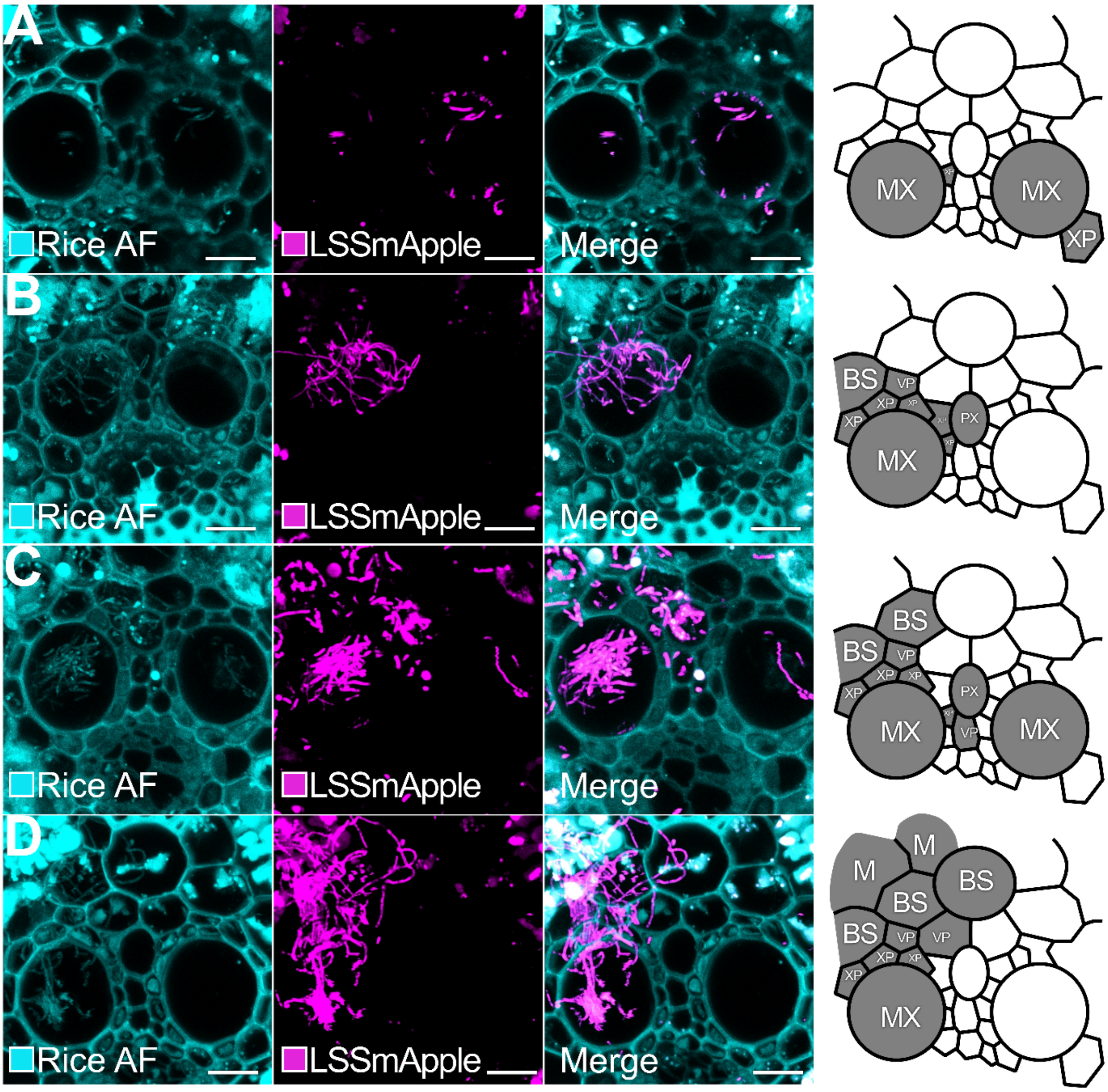


**Figure S8. Colonization pattern of Xoo in cross sections of major veins visualized by CLSM.** Representative images of filamentous Xoo observed in leaves of three independent experiments with 4 individual plants each. Apparent dispersal pattern of filamentous Xoo (grey) indicated in right column. **A)** Filamentous Xoo observed in metaxylem, xylem parenchyma, bundle sheath cells and mesophyll tissue. Individual filaments appear to reach lengths of 35 µm. Biological replicate 2 from independent experimental repeat 1. **B)** Filamentous Xoo >10 µm in the two metaxylem vessels. Apparent rod-shaped Xoo observed in xylem tracheid and xylem parenchyma cells. Biological replicate 1 from independent experimental repeat 2. **C)** Individual filaments of Xoo appear to reach lengths of 30 µm (observed here in 2D). Filaments traversed from metaxylem vessels to xylem parenchyma and bundle sheath cells. Biological replicate 3 from independent experimental repeat 3. **D)** Short filamentous Xoo (< 7 µm) observed in metaxylem and parenchyma cells. Biological replicate 3 from independent experimental repeat 3 (same independent repeat as in in Fig. S8C). Major vein cell pattern for cartoon was derived from S8D. MX: metaxylem vessel, BS: Bundle sheath cell, VP: vascular parenchyma cell, XP: xylem parenchyma cell M: mesophyll. Scale bar: 10 µm.

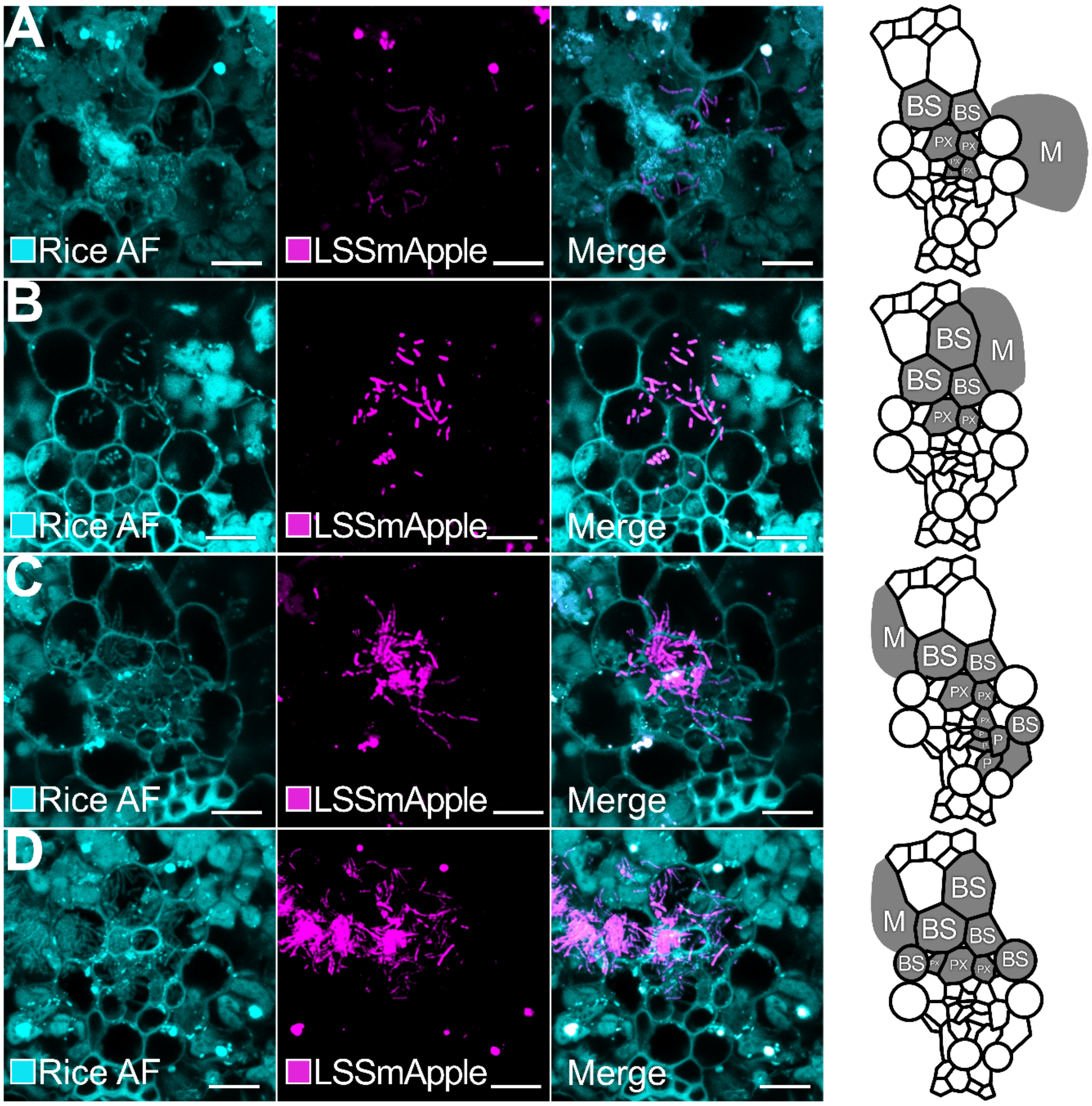


**Figure S9. Colonization pattern of Xoo in cross sections of minor veins visualized by CLSM.** Representative images of filamentous Xoo observed in leaves of three independent experiments with 4 individual plants each. Apparent dispersal pattern of filamentous Xoo (grey gradient) indicated in the cartoon (rightmost column). **A)** Short filamentous Xoo observed in protoxylem, bundle sheath and mesophyll tissue (approximately 6-8 µm). Other Xoo cells appeared rod-shaped in protoxylem and bundle sheath cells. Biological replicate 1 from independent experimental repeat 1.

**B)** Filamentous Xoo breach from protoxylem to bundle sheath and mesophyll tissue. High density of filamentous Xoo cells appear to cluster in parallel in bundle sheath cells and mesophyll tissue. Filamentous are approximately between 10-16 µm long. Biological replicate 1 from independent experimental repeat 2.

**C)** Individual filaments of Xoo appear to be up to 12.4 µm long. Filaments traverse from protoxylem to bundle sheath cells. Elongated Xoo cells appear to be septated. Biological replicate 3 from independent repeat 2 (same independent repeat as in Figure S9B).

**D)** Filamentous Xoo (approximately up to 21.2 µm long) observed in protoxylem and bundle sheath cells. Xoo cells appear to be septated. Biological replicate 4 from independent experimental repeat 3.

Minor vein cell pattern for cartoon was derived from S9B) and C. BS: Bundle sheath cell, PX: protoxylem (xylem parenchyma and tracheids), P: phloem, M: mesophyll.  Scale bar: 10 µm.

**Table S1. Xoo strains used in this study.** All strains were retrieved from the -80 °C glycerol stocks.

| **Strain** | **TALe** | **Targeted EBE sequence** | **Targeted *SWEET*** |
|---|---|---|---|
| $PXO99^{A}$ | PthXo1 | GCATCTCCCCCTACTGTACACCAC | *SWEET11a* |
| $PXO99^{A}$ | PthXo1 | GCATCTCCCCCTACTGTACACCAC | *SWEET11a* |
| ME2 | - | | non-virulent |

**Table S2. Rice lines used in this study.** All lines were retrieved from the rice seed stocks and tested for Hygromycin B resistance.

| **Name** | **Transformation Event #** | **Characteristics** | **Generation** |
|---|---|---|---|
| B01069 | [8-3] | pSWEET11a:g*SWEET11a*-GUSplus | T4 |
| B01075 | [10-2] | pSWEET11a:g*SWEET11a*-GUSplus | T4 |

**Table S3. Recipes for solutions for GUS histochemistry**

**Washing buffer**

| Component | Final concentration | For 50 ml |
|---|---|---|
| 0.5 M EDTA | 10 mM | 1 ml |
| 100 mM phosphate buffer pH 7 | 50 mM | 25 ml |
| 10 % triton X-100 | 0.1 % | 0.5 ml |
| 50 mM potassium ferrocyanide | 1 mM | 1 ml |
| 50 mM potassium ferricyanide | 1 mM | 1 ml |
| methanol | 20 % | 10 ml |
| sterile water | | Ad 50 ml |

**X-Gluc staining buffer**

| Component | Final concentration | For 50 ml |
|---|---|---|
| 0.5 M EDTA | 10 mM | 1 ml |
| 100 mM phosphate buffer pH 7 | 50 mM | 25 ml |
| 10 % Triton X-100 | 0.1 % | 0.5 ml |
| 50 mM potassium ferrocyanide | 1 mM | 1 ml |
| 50 mM potassium ferricyanide | 1 mM | 1 ml |
| methanol | 20 % | 10 ml |
| 100 mM X-Gluc | 2 mM | 1 ml |
| sterile water | | Add 50 ml |

**100 mM phosphate buffer, pH 7**

| Component | Volume |
|---|---|
| 0.5 M sodium phosphate dibasic ($Na_2HPO_4$) | 3 ml |
| 1 M sodium phosphate monobasic ($NaH_2PO_4$) | 1 ml |

**100 mM X-Gluc solution**

| Component | Final concentration | For 50 ml staining solution |
|---|---|---|
| X-Gluc | 100 mM | 50 mg |
| DMSO | | 1 ml |
